# Astrocyte-mediated immune shielding protects melanoma brain metastases from microglia/macrophage induced Interferon stress and metabolic collapse

**DOI:** 10.64898/2026.11.22.767511

**Authors:** A Herrero, Y Chen, FN Soria, E Armendariz-Burgoa, Z Xiong, T Ha, MD Boyano, Q Nguyen, B Arteta, A Benedicto

**Affiliations:** Cell Biology and Histology Department, School of Medicine and Nursing, University of the Basque Country, UPV/EHU, Leioa, Spain; Institute for Molecular Bioscience, The University of Queensland, Brisbane, Queensland, Australia; QIMR Berghofer, Brisbane, Queensland, Australia; IKERBASQUE, Basque Foundation for Science, Bilbao, Spain; Achucarro Basque Center for Neuroscience, Leioa, Spain; Vin University, Hanoi, Vietnam; Basque Center for Integrative Environmental and Human Health (EHU-iBio), María Goyri building, Barrio Sarriena s/n, 48940, Leioa, Spain; Biocruces Health Research Institute; Instituto Biofisika (UPV/EHU, CSIC), University of the Basque Country, Leioa, 48940, Spain; Department of Biochemistry and Molecular Biology, University of the Basque Country, UPV/EHU, Leioa, Spain; IrsiCaixa, 08916, Badalona, Barcelona, Spain

## Abstract

Brain metastasis severely limits melanoma prognosis driven by permissive brain tumor microenvironment (TME), wherein astrocytes actively favor progression. Yet the microenvironmental consequences of disrupting tumor-reactive proliferating astrocytes (TARA) remain uncharacterized. Using transgenic mouse model, we selectively eliminated TARA in intracranial melanoma. Immunohistochemistry and spatial transcriptomics uncovered that TARA are essential for early and late tumor growth; notably, TARA depletion reduced 34-fold late melanoma growth. This depletion triggered proinflammatory, antigen presenting microglial/macrophage infiltration leading to melanoma cell tumorostatic reaction. Mechanistically, TARA ablation abrogated melanoma cell cycle machinery, upregulated interferon response and p53 pathways, resulting in stress-mediated, cytostatic and hypertrophic phenotype. Spatial transcriptome identified TARA-melanoma interaction through Lgals1-Itgb1, Vim-Cd44, reduced following TARA depletion. Moreover, increased Mif-Cd74 and Apoe-Scarb1 signaling in surviving cancer cells suggests potential activation feedback loops as a last resort to evade immune clearance. Targeting TARA rewires the metastatic TME, unleashing antitumor immunity to suppress brain metastasis.

**Graphical abstract:** 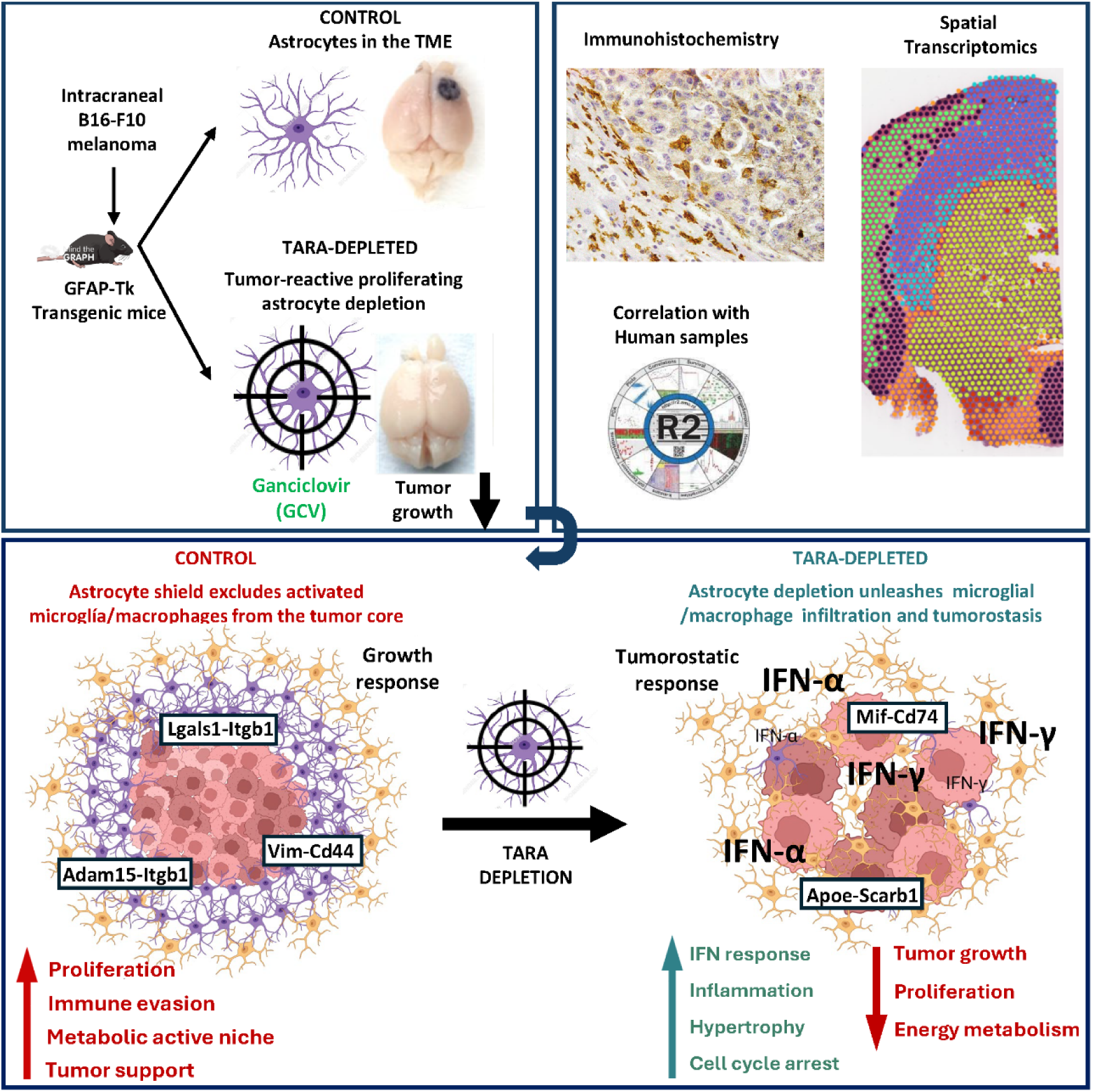

## Introduction

Melanoma incidence is rising every year. While the primary lesion is usually resectable with 5-year disease free survival of 98 %, the dissemination of the primary tumor to other organs remains as the most challenging scenario when dealing with this aggressive skin cancer. Melanoma tends to metastasize to lymph nodes, lungs, brain and liver, which reduces the 5-year overall survival notably, with only 35 % of the patients achieving this timeframe. Among these metastatic sites, the central nervous system (CNS) represents one of the most lethal locations due to the restrictive nature of the blood brain barrier (BBB) and the protective, tumor-promoting niches established by resident cells (1).

Once melanoma cells extravasate into the brain parenchyma, they must interact with resident cells to create a favorable tumor microenvironment (TME) to proliferate and survive (2). Astrocytes represent the most abundant glial cell in the brain, and are responsible for maintaining organ homeostasis, metabolic support and BBB maintenance under normal physiological conditions (3). In fact, in recent years astrocytes have arisen as supporters of brain cancer and brain metastasis through several mechanisms. Astrocytes are known to boost cancer cell proliferation through the secretion of neurotropic factors and pro-inflammatory cytokines (Burn et al., 2021), while blocking the cytotoxic response of lymphocytes towards tumor cells (4). Astrocytes mediate glioma and breast and lung metastatic cell invasion and resistance to therapy through conexin 43 /Cx43) protein (5,6) while they are found in the invasive front of medulloblastoma, linked to aggressive cancer phenotype and therapy resistance (7). Besides, peritumoral astrocytes expressing GFAP and platelet-derived growth factor receptor α (PDGFRα) are associated with worse prognosis in glioma (8), pointing out astrocytes as relevant mediators of tumor growth in the brain.

In the specific context of melanoma brain metastasis (MBM), this bidirectional crosstalk is heavily hijacked by colonizing cells to fuel their invasive and survival capacities (9). Upon parenchymal infiltration, melanoma cells trigger a localized pro-inflammatory activation in surrounding astrocytes, which respond via upregulation of CXCL10 (10). Furthermore, this interplay prompts a phenotypic transformation driven by astrocytic STAT3 signaling pathways, which correlates with reduced survival in patients (11). This reactive state or astrogliosis, alters the local secretome, providing metabolic support and creating an immunologically “cold” TME that allows metastatic cells to evade local immunosurveillance (11). Consequently, understanding the functional heterogeneity of these responsive glial population is crucial to disrupting the niche they provide to aggressive melanoma cells.

Despite the recent advances between astrocytes-tumor interplay, there is still an unmet need to understand the astrocyte dependency of cancer cells in the brain and how astrocytes modulate both cancer cells and brain TME conforming different cell populations. To address the relevance of proliferating tumor-activated reactive astrocytes (TARA), we used an in vivo approach using the transgenic GFAP-Tk mice to deplete TARA from melanoma brain tumor growth. Using this model, only tumor-reactive proliferating astrocytes undergo apoptosis, while the rest of this pericyte-like cell population remains unaffected. This specificity allows us to characterize the relevance of TARA during early and late tumor growth in the brain.

## RESULTS

### TARA drive both early and late melanoma progression within a protective brain niche

Astrocytes proliferate upon brain damage, in a process called astrogliosis. Because tumor progression mimics a non-healing wound, reactive astrocytes tend to accumulate in the peritumoral niche reminiscent of a glial scar (Figure 1A). To interrogate the functional dependency of melanoma on these tumor-associated reactive proliferating astrocytes (TARA), we utilized the GFAP-Tk transgenic mouse model. This system allows for the selective Ganciclovir (GCV)-mediated depletion of proliferating astrocytes-the specific population engaged in direct crosstalk with invading melanoma cells-while not affecting homeostatic counterparts. Following intracranial injection of B16-F10 melanoma cells, mice were treated with either saline (Control) or GCV (TARA-depleted) from day 4 to day 11 and analyzed at day 12 (Figure 1B). Immunohistochemical evaluation for Ki67 and GFAP revealed that astrocytes proliferate to maintain the forming barrier as the tumor evolves, as reported in Figure 1C. TARA depletion diminished Ki67 positive astrocytes almost completely from the periphery of the melanoma tumor mass. Moreover, the astrocytic scar was missing when TARA were depleted. Crucially, non-proliferating astrocytes, located within the immediate parenchymal vicinity of the nascent foci (<100µm), remained unaffected, retaining their characteristic star-like shape and demonstrating the high specificity of this system (Figure 1C). GCV treatment exerted no confounding effects on wild type littermates (Suppl. Figure 1A, B).

**Figure 1.**
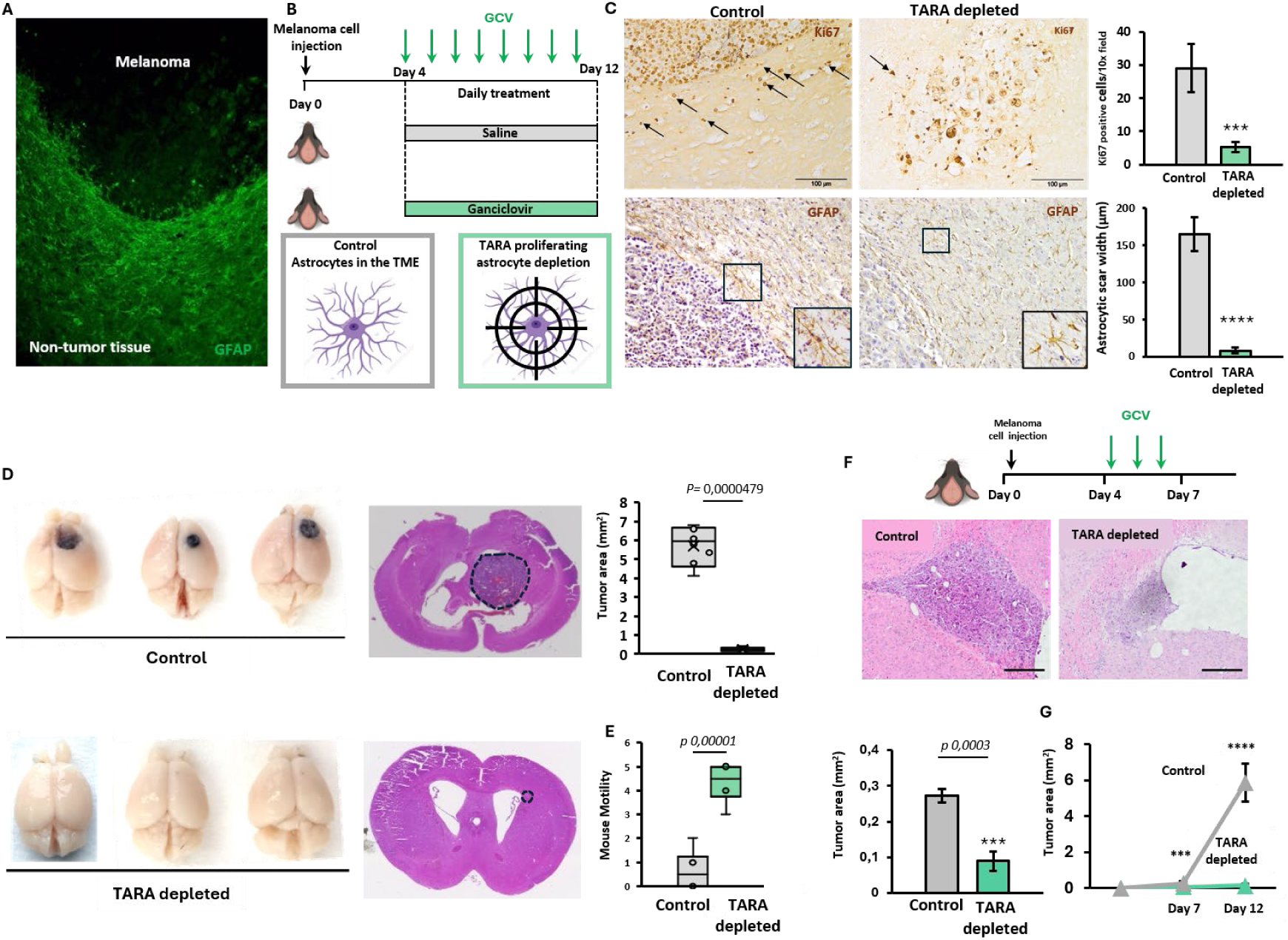
Histology analysis shows TARA depletion related tumor growth reduction in mice. A) Image from GFAP immunofluorescence (green) in melanoma tumor bearing mouse brain. B) TARA depletion model using GCV from day 4 to 11 followed by sacrifice. C) On the left, images from Ki67 and GFAP immunohistochemistry in control and TARA- depleted brain with quantification in the right bar plots. Scale bar: 100 µm. D) On the left, representative images of the macroscopic tumors in control (upper panel) and TARA-depleted (bottom panel) brains at 12 days; On the right, hematoxylin-eosin (H-E) staining from representative images of microscopic tumors at 12 days (n=6/group). E) On top, tumor area measurement between control and TARA-depleted mice; on the bottom, motility assay between control and TARA-depleted mice. F) In vivo early tumor growth TARA depletion for 7 days model. Middle panels show H-E staining of the brain tissue after 7 days of experimental tumor growth; on the bottom left tumor area measurement comparing control and TARA-depleted (n=3/group); and on the right tumor growth follow up using both time points. Scale bar: 50 µm. Differences between control group and TARA-depleted group were considered statistically significant for **p<0.01, ***p<0.001, ****p<0.0001.

We next explored whether blocking this astrocytic response impacts melanoma progression. TARA ablation rendered a 95 % reduced melanoma burden at day 12compared to the control group (Figure 1D). While control group exhibited an average of 5,8 mm^2^ tumor area, TARA-depleted mice showed 34-fold reduced tumor growth, with lesions averaging just 0,17 mm^2^ (Figure 1D, Suppl. Figure 1A). This severe tumor restriction directly translated into preserved quality of life (QoL) and neurological integrity. While control group developed severe locomotor and coordination deficits, moving erratically by day 12, TARA-depleted mice remained symptom free, maintaining physical dynamics, as quantified using the Experimental Autoimmune Encephalomyelitis (EAE) neurological scoring scale, which ranges from 0 to 5 (or up to 6 in the more severe cases) to grade the progression of paralysis in animal models (Figure 1E).

To define the kinetics of this dependency, we next characterized the early brain response to melanoma growth by analyzing tumor development at day 7, after 3 days of GCV treatment (Figure 1 F). Even at this early checkpoint, TARA elimination significantly restricted tumor progression up to 4-fold (0.27 mm2 in control vs 0,08 mm2 in TARA-depleted), indicating that astrocytes support melanoma growth from its initial stages. Longitudinal analysis between day 7 and 12 revealed an exponential, greater than 20-fold increase in tumor area in control mice, whereas TARA-depleted tumors expanded 2-fold (Figure 1G). Together these temporal windows demonstrate that tumor-reactive proliferating astrocytes constitute a required microenvironmental catalyst to sustain exponential growth phase of intracranial melanoma.

### TARA elimination triggers sustained cytostatic and hypertrophic phenotypes

To uncover the mechanisms driving to the dramatic tumor restriction, we next analyzed the histopathological features of remaining melanoma cells upon TARA elimination. Morphometric analysis revealed striking phenotypic alterations in cancer cells, characterized by cellular hypertrophy and 3-fold increase in nuclear size at both early (7 days) and late (12 days) timepoints (Figure 2A-B). Because cell size enlargement coupled with cell growth inhibition account for cell cycle arrest and given that astrocyte-derived factors promote cancer cell proliferation (12,13), we examined tumor replicative dynamics through Ki67 immunohistochemistry. We confirmed cell cycle defects within B16-F10 cells in TARA-depleted tumors, revealing 50% reduction in Ki67 signal intensity at day 7, showing the same trend at day 12, with reduced Ki67 cell number and intensity (Figure 2A and 2B). Given the significant role of TME in immune evasion, we hypothesized that TARA elimination might additionally accelerate apoptotic tumor death. However, cleaved Caspase-3 profiling at both timepoints revealed negligible apoptosis within the core of TARA-depleted tumors, while the scarce Caspase-3 signal was constrained to the peritumoral border, reflecting GCV-mediated astrocyte ablation rather than direct tumor cell apoptosis (Figure 2A and 2B).

**Figure 2:**
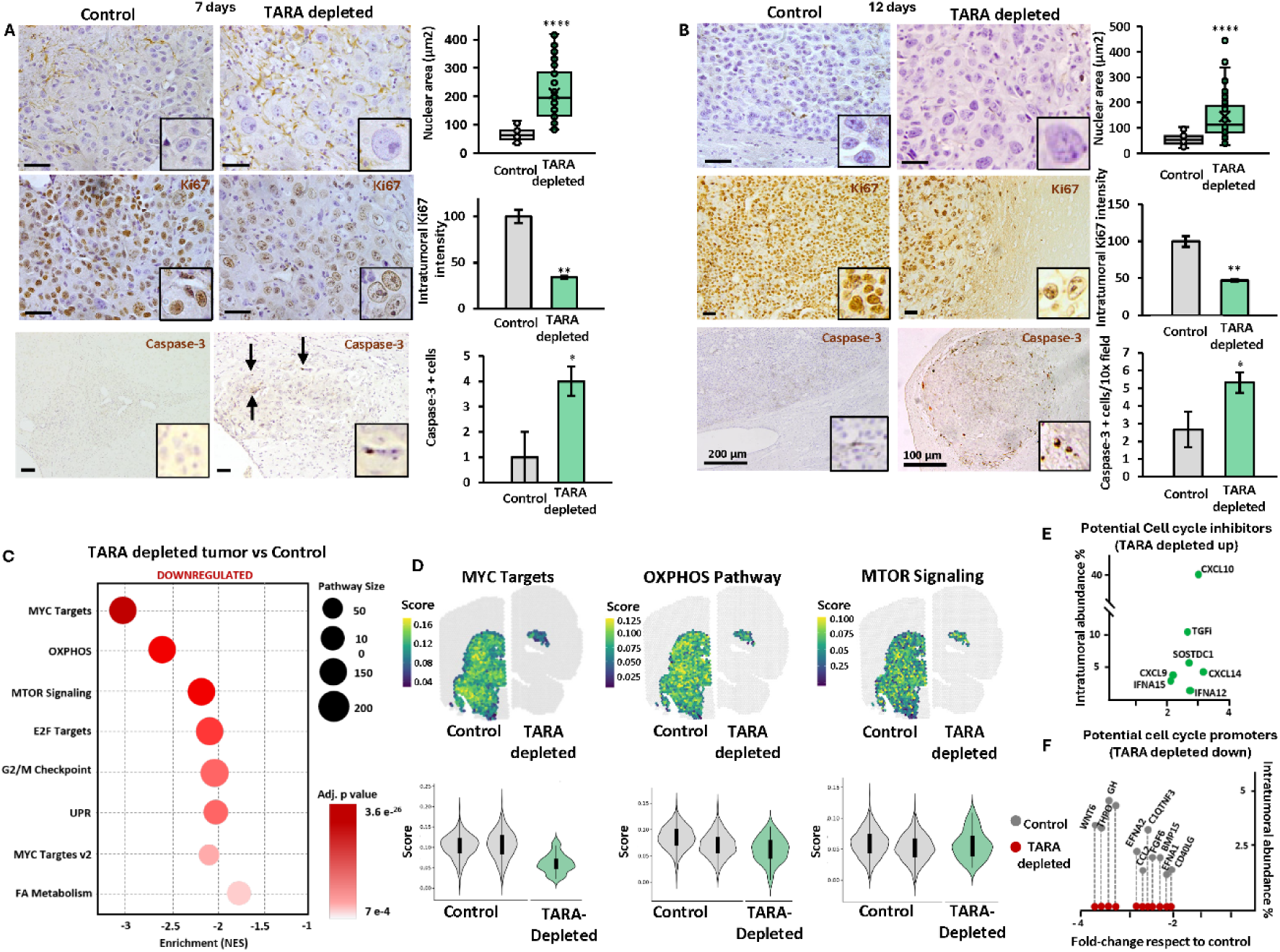
Characterization of melanoma tumors reveals impaired cell cycle. A) Cell and nuclear size analysis, Ki67 and Caspase-3 immunohistochemistry during early tumor growth (7 days) in control and TARA-depleted mice. Scale bar: 50 µm. B) Cell and nuclear size analysis, Ki67 and Caspase-3 immunohistochemistry during late tumor growth (12 days) in control and TARA-depleted mice. Scale bar: 50 µm as otherwise stated. C) Gene Ontology showing reduced pathways in TARA-depleted tumors compared to control tumors at late tumor growth (12 days). D) Spatial mapping of top three downregulated pathways, MYC targets, OXPHOS and MTOR Signaling in control and TARA-depleted mouse tumor. E) Intratumoral abundance of potential cell cycle inhibitors upregulated in TARA-depleted tumors. F) Intratumoral abundance of cell cycle promoters downregulated in TARA-depleted tumors compared to control tumor. Differences between control group and TARA-depleted group were considered statistically significant for *p< 0.05, **p<0.01, ***p<0.001, ****p<0.00001.

To map the transcriptional programs driving this non-apoptotic, antiproliferative and hypertrophic phenotype, we carried out spatial transcriptomics. Pathway enrichment and spatial mapping analyses revealed a robust remodeling of the melanoma transcriptome that closely mirrored out histological observation. TARA-depleted tumor exhibited a profound downregulation of essential cell cycle machinery, including MYC Targets (-3.15 NES), E2F targets (-2.2 NES) and G2/M targets (-2.15 NES) (Figure 4C). Furthermore, TARA depletion severely suppressed Oxidative Phosphorylation (-2.61 NES) and Fatty Acid metabolism (-1.76 NES) programs, suggesting a critical loss of astrocytic metabolic support. Linking both proliferative and metabolic blocks, MTOR Signaling pathway was markedly downregulated (-2.23 NES) and, accompanied by impaired Unfolder Protein Response (UPR) (-2,14 NES), consistent with low-rate, stress-induced metabolic state (Figure 4C). Cell deconvolution analysis further supported this tumorostatic collapse, revealing a pronounced contraction of the melanoma transcriptomic signature within TARA-depleted niche, accompanied by profound diminishment of proliferating astrocyte population, further confirming previous results (Suppl. Figure 2A-B).

To further infer if intercellular communication could play a role in the observed poor melanoma-supportive microenvironment, we further analyzed intercellular communication networks. Spatial expression analysis revealed potential antiproliferative mediators including Ifna12 and Ifna15, were upregulated over 4-fold in TARA-depleted tumor (Figure 2E). Conversely, potential oncogenic and mitogenic stimuli, including Wnt6 and Fgf6, were diminished from the TME upon TARA depletion (Figure 2F), which may also account for reported melanoma cell growth impairment.

At a whole, these histopathological and spatial transcriptomic data suggest that TARA depletion drives an energetic and proliferative collapse in melanoma, forcing cancer cells into a tumorostatic hypertrophic survival state independent from apoptosis.

### TARA elimination suppressed protumoral niche while rewiring brain towards a hostile microenvironment

We have shown that TARA elimination impairs melanoma growth in the brain. To characterize the peritumoral area upon TARA depletion and put some light on the reported tumor restriction, we next characterized how preventing the establishment of this protumoral niche alters the peritumoral architecture. We estimated the cell composition of the striatal peritumoral niche (within two Visium spots of the tumor) by deconvolution scaled to the nuclei counted in each spot. This approach revealed reduced reactive astrocyte transcripts in the area surrounding TARA-depleted tumor (Figure 3B) Among non-tumor cells, astrocytes made up 17.5% and 18.4% of the control peritumoral niche but only 12.9% when TARA recruitment was blocked, below the 95% confidence intervals of both controls (Figure 3C). Microglia and other immune cells took up a larger share in TARA-deleted peritumoral area (microglia 33.3% vs 26.6% and 29.8%; other immune cells 14.3% vs 11.3% and 10.0%). OPC and oligodendrocyte (5.1% vs 6.1% and 6.8%) and vascular (7.6% vs 7.8% and 9.0%) shares were close to control. The neuron share did not recover (17.8% vs 24.4% and 20.5%; Figure 3C). Thus, preventing TARA-driven niche formation abolishes the astrocyte-rich, melanoma-infiltrated border, and immune cells rather than neurons fill it.

**Figure 3.**
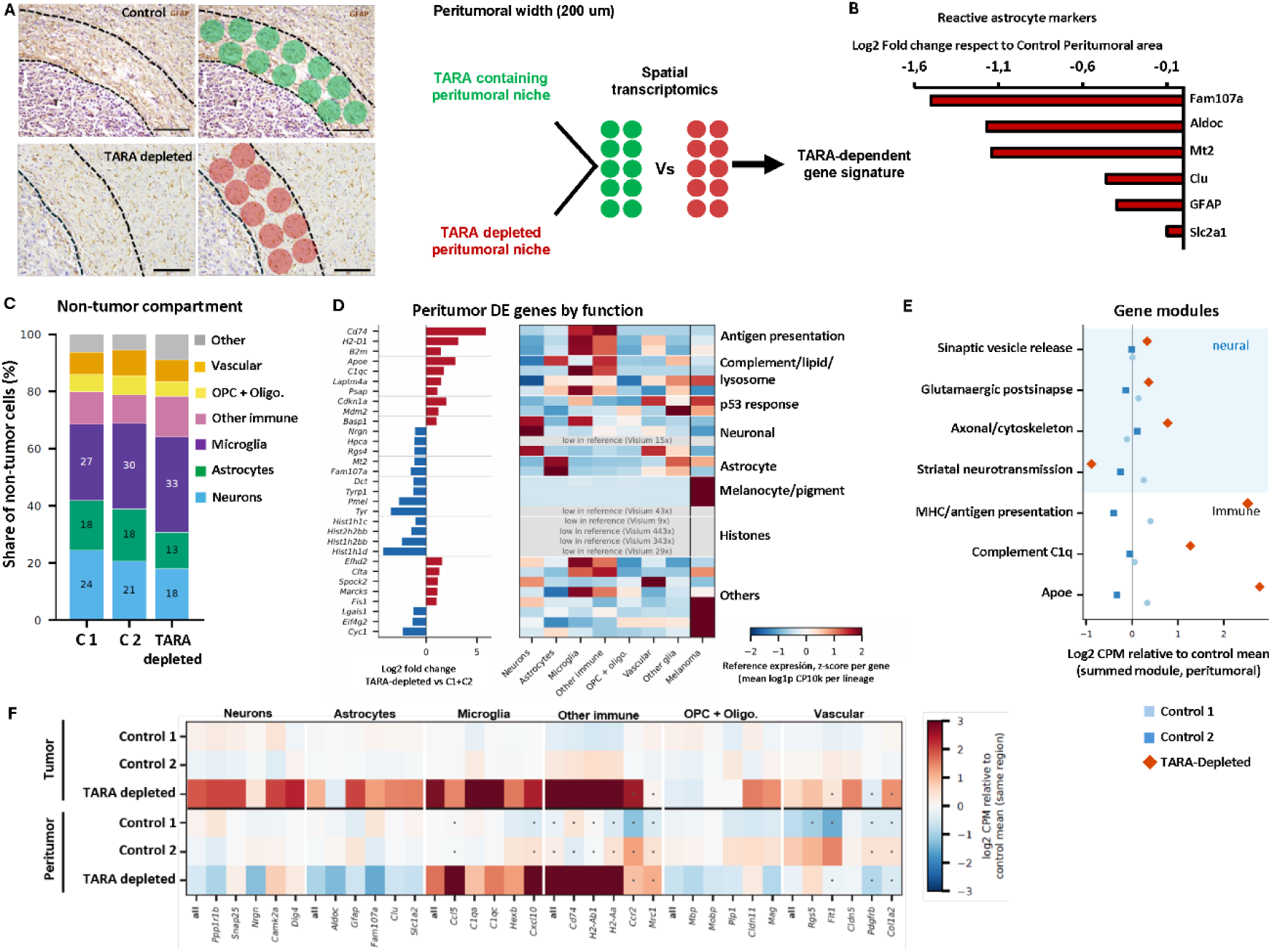
TARA depletion prevents the astrocyte-rich peritumoral niche and engages an antigen-presenting innate immune border. A) GFAP immunohistochemistry showing spatial mapping of compared peritumoral area and subsequent data analysis in control (Green) and TARA-depleted (red) tumors B) Reduced reactive astrocyte markers found in TARA-depleted peritumoral area compared to control peritumoral area. (C) Share of non-tumor cells per lineage in the peritumoral niche and in distant striatum of the same section (≥ 3 spots from the tumor). D) Genes differentially expressed in the peritumoral niche, TARA-Depleted vs controls (edgeR quasi-likelihood on pseudobulk spot pools; 328 genes tested; FDR < 0.05, |log2FC| ≥ 1), grouped by function, with their expression across lineages in the single-cell reference (scaled to each gene’s maximum). (E) Neural and innate immune gene modules in the peritumoral niche: summed-module log2 CPM relative to the mean of the two controls; one point per animal. (F) Lineage marker genes in the tumor and peritumoral niche, log2 CPM relative to the control mean in the same region; “all” is the summed marker set; dots mark values based on fewer than 5 transcripts. One section per animal; comparisons are descriptive.

To determine the functional landscape of the peritumoral area, we organized the differentially expressed transcripts into functional groups (Figure 3D) and scored pre-specified neural and immune gene modules (Figure 3E). Transcripts governing synaptic vesicle release (log2 +0.33 relative to the control mean), glutamatergic postsynaptic function (+0.36) and axonal maintenance (+0.78) were maintained at the TARA-depleted border, whereas striatal medium-spiny-neuron transcripts were lower (−0.89; Ppp1r1b, Nrgn, Hpca). This indicates that neural transcriptional activity is not suppressed at the border when the TARA-derived scar is prevented. Crucially, the peritumoral environment remained hostile to melanoma progression. Immune response genes of microglia and macrophages displayed 1-to 6-fold (log2) upregulation at the TARA-depleted border: Cd74 (log2FC 5.9), H2-D1 (3.1), Apoe (2.9), C1qc (1.8) and B2m (1.4) (Figure 3D). Consistently, the MHC / antigen-presentation (+2.5), complement C1q (+1.3) and Apoe (+2.8) modules rose (Figure 7E). Lineage marker genes confirmed lower astrocyte markers (Aldoc, Fam107a) and higher microglial (C1qa, Cxcl10, Ccl5) and antigen-presenting (Cd74, H2-Aa, H2-Ab1) markers at the TARA-Depleted border (Figure 3F). This indicates that, in the absence of TARA-mediated immunosuppression, local innate immunity remains actively engaged to suppress tumor progression.

### TARA depletion unleashes robust intratumoral interferon and inflammation response

Given the dynamic intercellular interactions within the tumor microenvironment, the loss of a constituent cell type during melanoma progression would be expected not only to diminish specific responses, but also to elicit compensatory or emergent changes among the remaining populations. Moreover, previous findings suggest local immune response towards melanoma. To characterize such alterations, we analyzed transcriptional programs upregulated upon TARA depletion. Spatial transcriptomics revealed a robust activation of inflammatory programs, highlighted by 2.5-2.7-fold enrichment in Interferon (IFN) Alpha/Gamma Responses and General Inflammation pathways compared to control tumors (Figure 4A). Furthermore, specific immune-activating cascades, including Complement and IL2-STAT5 signaling pathways exhibited 1.9 and 1.7-fold enrichments, respectively, alongside with a 2-fold enrichment in the Allograft Rejection Pathway (Figure 4A). Concurrently, p53 Pathway, intrinsically linked to suppressed cell cycle progression was upregulated (1.7-fold). Interestingly, KRAS Signaling UP Pathway was enriched 1.9-fold, which may account for cell senescence or immune related inflammation (Figure 4A). Spatial mapping of the core enrichment genes driving these dominant inflammatory pathways showed a robust upregulation localized across both intratumoral core and the peritumoral border, suggesting a coordinated response driven by both cancer cells and infiltrating immune cells (Figure 4B).

**Figure 4.**
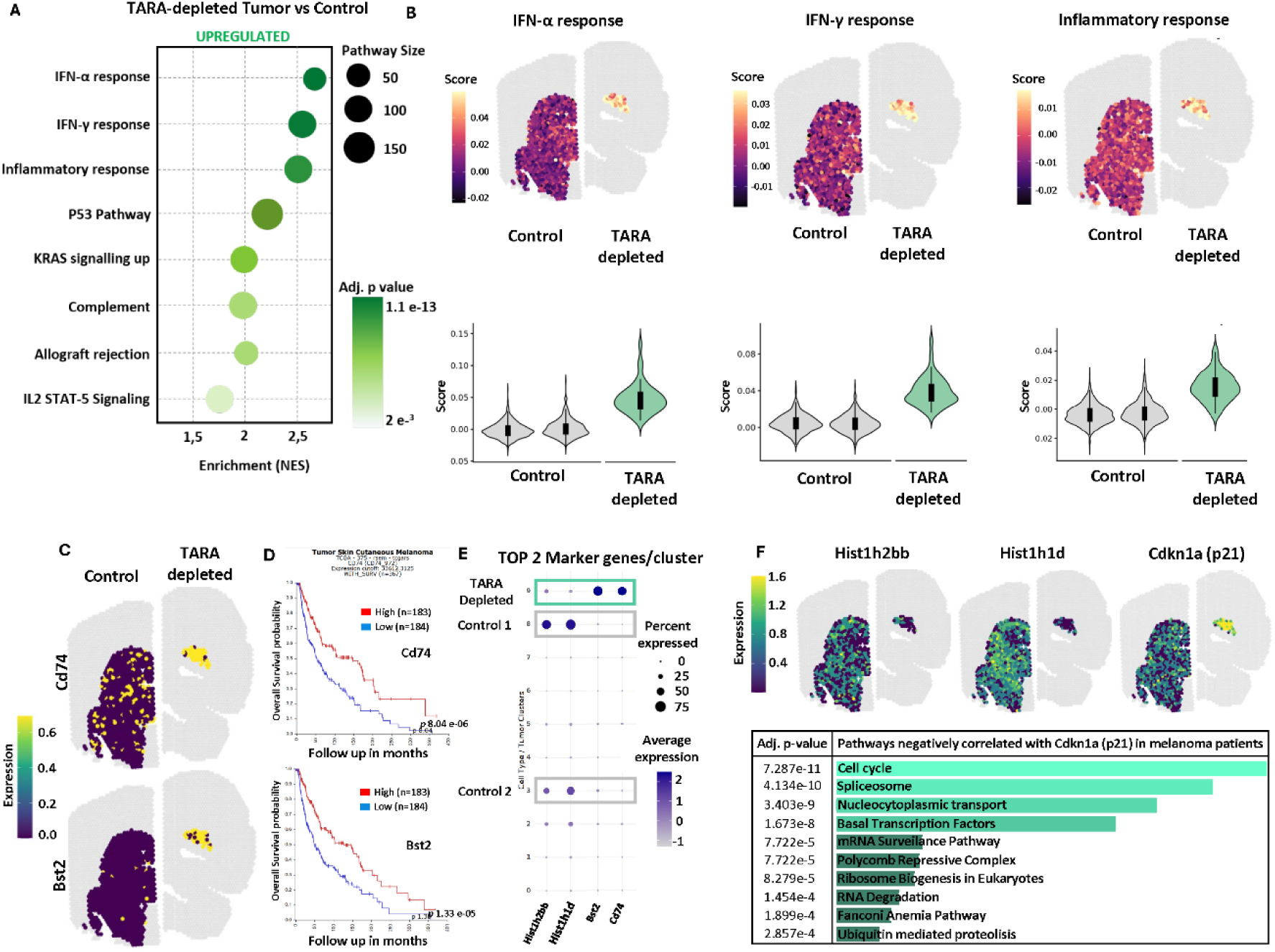
Upregulated pathways in melanoma tumors reveal strong inflammatory response. Upregulated Gene Ontology showing upregulated pathways in TARA-depleted tumors compared to control tumors at late tumor growth (12 days). B) Spatial mapping genes of top three upregulated pathways, IFN-α response, IFN-γ response and Inflammatory response in TARA-depleted tumors compared to control tumor. C) Spatial mapping of Cd74 and Bst2 genes in control and TARA-depleted tumors. D) Kaplan-Meier survival curves in 375 melanoma patient cohort showing CD74 and Bst2 related to improved overall survival. E) Dot plot showing the most differentially expressed genes between clusters, key genes in cell cycle dynamics and in interferon response. F) Spatial mapping of Hist1h2bb, Hist1h1d and Cdkn1a (p21) in control and TARA-depleted tumors with negatively correlated pathways with p21 in 375 melanoma patient’s cohort.

Consistent with this spatial distribution, we mapped the expression of Cd74 and Bst2, the top two markers’ genes for TARA-depleted tumor-associated clusters, both tightly linked with interferon-responsive programs. Spatial profiling revealed an overlapping pattern for both transcripts extending across tumor boundary into the adjacent parenchyma (Figure 4C–D). To establish the clinical relevance of this transcriptional signature, we examined the association of CD74 and BST2 expression with clinical outcomes in a public cohort of 375 melanoma patients. Elevated expression of both genes was significantly associated with prolonged overall survival, (p values of 8.04 × 10⁻⁶ and 1.33 × 10⁻⁵, respectively; Figure 4D), indicating that the transcriptional features emerging after TARA depletion are consistent with favorable prognostic signatures in human melanoma.

Conversely, the top two control tumor cluster markers, Hist1H2bb and Hist1h1d, were both almost absent in spatial mapping of TARA-depleted mice tumors, confirming the collapse of the tumor replication machinery (Figure 4F). Parallel to this, p53 pathway, commonly activated in response to interferon mediated stress (14), was upregulated in TARA-depleted tumors. Downstream of p53, the canonical cyclin-dependent kinase inhibitor p21 (Cdkn1a) acts as a critical cell cycle brake. Spatial mapping of p21 confirmed a robust upregulation of Cdkn1a expression in TARA-depleted tumors (Figure 4F). This induction, along with the profound loss of Hist1hbb and Hist1h1d expression (Figure 4F), underlies the observed proliferative arrest, suggesting IFN and Inflammation-driven tumorostatic reaction leading to constrained melanoma growth. To validate this mechanism in human melanoma, we analyzed genes negatively correlated with CDKN1A (p21) in publicly available human melanoma datasets. Pathway analysis revealed that cell cycle progression is the most significantly repressed program (*p* 7.287e^-11^), alongside essential cellular machinery such as Spliceosome pathway (*p* 4.134e^-10^), Nucleocytoplasmic transport (*p* 3.403e^-9^) and Basal Transcription Factors pathways (*p* 1.673e^-8^) (Figure 4F). Together these findings suggest that preventing TARA-mediated peritumoral niche establishment unleashes a localized, interferon mediated tumorostatic response that mirrors favorable prognosis in patients.

### Peritumoral TARA depletion favors the infiltration of activated iba-1^+^ cells into the tumor core

Astrocytes create a protective shield for tumor cells to grow (15,16). By selectively eliminating peritumoral TARA, we hypothesized that this physical barrier is dismantled, altering the immune landscape of the niche. Given the robust inflammatory and Interferon response observed in TARA-deficient tumors and the peritumoral niche composition, we next investigated whether this phenomenon was driven by macrophages and main source of inflammatory stimuli in the brain, microglia. Immunohistochemical profiling of Iba-1 revealed a highly distinct spatial distribution of these cells. In control tumors Iba-1^+^ cells tend to accumulate densely in the peritumoral niche, seemingly sequestered within the astrocytic scar, with slight infiltration into the tumor mass (Figure 5A). Remarkably, TARA ablation completely reshaped this peritumoral niche; Iba-1^+^ cells extensively infiltrated the tumor foci, penetrating deep into the melanoma tumor the core (Figure 5B). Spatial distribution analysis confirmed that in the presence of peritumoral TARA, Iba-1 expression was mainly constrained to the outer tumor margins, with the tumor boundary acting as a limiting barrier for most cells (Figure 5C; Suppl. Figure 3). Indeed, Iba-1 accumulation pattern closely resembled that of GFAP^+^ astrocytes at the peritumoral space (Suppl. Figure 4A). Conversely, TARA depletion abolished this spatial restriction, unleashing a robust infiltration into the melanoma mass. While control tumors maintained a sharp expression gradient between the peritumoral border and the tumor core, TARA-depleted lesions exhibited uniform, high-density Iba-1 infiltration pattern across the entire tumor landscape (Figure 5D). Quantification of Iba-1^+^ area supported this observation, revealing a significant 5.5-fold increase in intratumoral area positive for Iba-1^+^ in TARA-depleted tumors, increasing from 4,8% in control tumors up to 27,9% in TARA-depleted tumors (*p*=0,0014; Suppl. Figure 3C). This extensive Iba-1^+^ cell influx was evident even at early timepoint (day 7), where most melanoma cells surrounded by Iba-1 positive cells and displayed 3-fold enlargement in nuclear and cellular size (Suppl. Figure 4B). This histopathological finding aligns perfectly with spatial mapping of activated microglia transcriptomic signature (detailed in Suppl. Figure 5); while nearly 100% of tumor spots positively scored for activated microglia, only 28.9 % of spots in control tumors localized with this signature (Figure 5E). Quantitatively, TARA depletion induced a 4-fold increase in the average microglial signature score (Figure 5E), pointing out the peritumoral astrocytes as a plausible physical barrier that impede microglial-tumor interaction. Cell deconvolution further supported this observation, revealing 3-fold increase in activated microglia within the core of TARA-depleted tumors (Suppl. Figure 2C).

**Figure 5.**
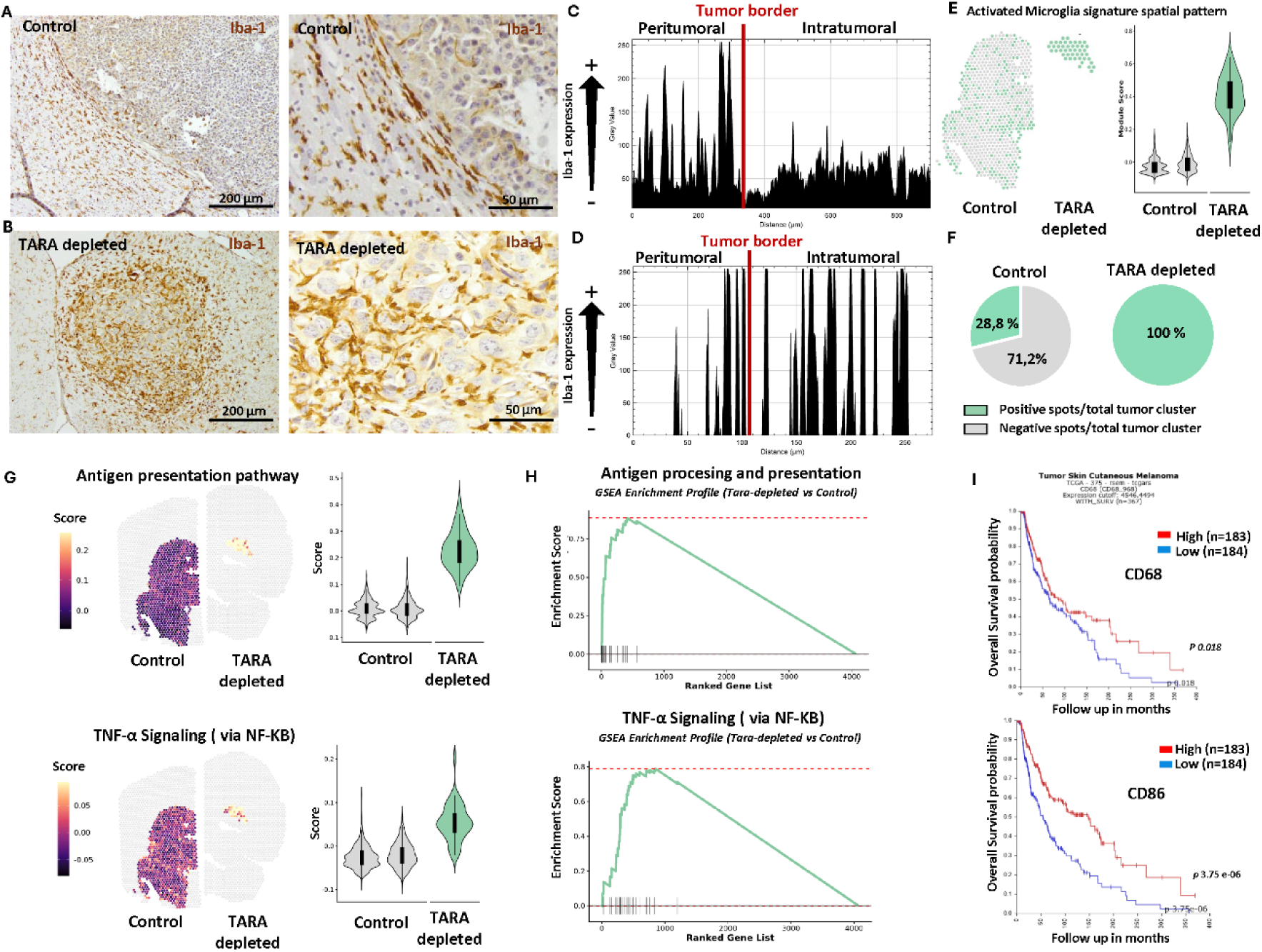
TARA elimination facilitates Iba-1^+^ cell infiltration and activation. A-B) Immunohistochemistry of Iba-1 in control (upper panel) and TARA-depleted brains (bottom panel). C-D) Analysis of Iba-1 expression in the peritumoral and intratumoral area in control (up) and TARA-depleted brains (bottom). E) Spatial mapping of microglial activation signature in control and TARA-depleted tumors (left) and violin plots showing average expression of microglial activated signature (right). F) Cake chart showing positive (green) and negative (grey) spot percentage for microglial activation signature in control and TARA-depleted tumors using spatial mapping. G) Spatial mapping of genes related with *Antigen presentation* and *TNF-α signaling through NFkb* pathway with correspondent violin plots and GSEA enrichment profiles in control and TARA-depleted tumor. I) Kaplan Meier survival curves from 375 melanoma patient cohort showing increased overall survival related to CD68 (upper panel) and CD86 (lower panel) expression.

To functionally characterize this infiltrating population, we spatially mapped two dominant microglial activation pathways; Antigen presentation and TNF-α Signaling via NF-Kβ. Both pathways exhibited a 3-fold enrichment in the intratumoral compartment, aligning precisely with the microglial infiltration pattern (Figure 5G-H). Finally, to evaluate the translational significance of these findings, we analyzed human melanoma dataset to assess the prognostic value of this inflammatory myeloid phenotype. Consistent with our murine data, elevated expression of both the activated microglial marker CD68 (*p*=0.018) and the pro-inflammatory/antigen presenting marker CD86 (*p*=3.75 e^-6^) significantly correlated with prolonged overall survival in a clinical cohort of 375 melanoma patients (Figure 5I). Taken together, these data demonstrate that TARA depletion dismantles the astrocytic shield, allowing activated, antigen-presenting microglia/macrophage to invade the melanoma mass, correlating with an interferon-mediated tumorostatic response that aligns with favorable patient outcome.

### TARA ablation rewires the metastatic TME by disrupting pro-tumorigenic signals and promoting melanoma-microglia/macrophage immunogenic interplay

To dissect the structural and communicative reorganization of the brain metastatic TME dependent on TARA depletion, we performed cell deconvolution and ligand-receptor interaction analysis using Visum 10x. Melanoma cells exhibited a dense network of protumorigenic crosstalk with astrocytes in the Striatum, that spatially correlated with the active tissue invasion into healthy brain parenchyma (Figure 6A). Remarkably, specific depletion of TARA completely dismantled these pro-invasive signals. We observed a total rewiring of the astrocyte-melanoma communication network in the peritumoral area, with loss of several ligand-receptor interactions, such as Vimentin-Cd44, Lgals1-Itgb1 or Adam15-Itgb1 (Figure 6C; Suppl. Figure 6). Instead, the remaining hypertrophic tumor cells underwent a profound niche shift, increasing their local immunological engagement with the massive infiltrating inflammatory microglia/macrophages (Figure 6B). Spatial cell-cell communication modeling revealed two of the potential driver axes present in this interplay: MIF-CD74 stress survival loop and the Apoe-Scarb1 lipid-mediated metabolic support, alongside Vim-CD44 and Lgals1-Itgb1 (Figure 6D; Suppl. Figure 7). Interestingly, inflammatory microglia–tumor interactions were detected in only a small fraction of spots in control brain but were widespread across nearly all spots in TARA-depleted tumors (Figure 6B), consistent with histological evidence of increased intratumoral Iba-1⁺ cell infiltration. Together these spatial transcriptomic data suggest that eliminating TARA-conformed protective reactive peritumoral niche deprives melanoma of its invasive communicating network, forcing the tumor into isolated, defensive state defined by intense microglial/macrophage engagement and a complete arrest of progressive tissue invasion.

**Figure 6.**
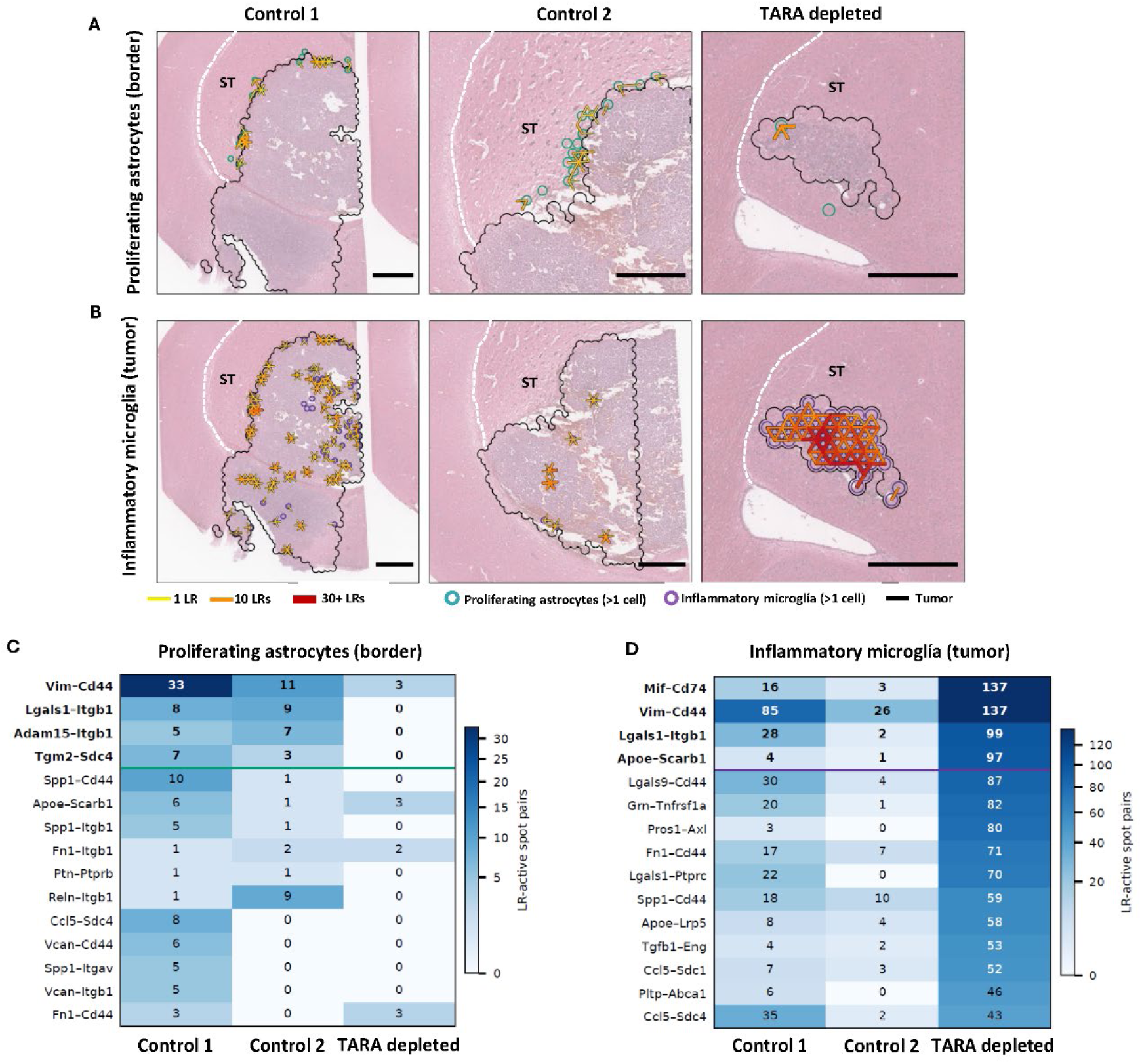
TARA depletion reshapes melanoma cell-astrocyte and melanoma-microglia/macrophage interaction. A) spatial mapping of striatal proliferating astrocytes (green dots) and melanoma interaction overlapping with the tumor border (black line). B) Spatial mapping of Melanoma-inflammatory microglia (purple circles) interplay in the intratumoral region. C) Ligand-receptor (LR) active spots occurring in the tumor border between proliferating astrocytes and melanoma cells. D) L-R active pairs taking place in the intratumoral regions between inflammatory microglia and melanoma. Scale bar: 500 µm. ST: Striatum.

## Discussion

The orchestration of the brain tumor microenvironment by stromal cell represents a key yet poorly understood step during metastatic colonization of the brain. Our study describes that TARA orchestrate TME remodeling that sustains melanoma colonization from early seeding to advanced progression. Mechanistically, TARA depletion dismantles critical peritumoral interplay axes (including Lgals1-Itggb1 and Vim-Cd44), breaching the astrocytic barrier to allow a progressive and massive infiltration of antitumor Iba-1^+^ inflammatory myeloid cells forcing melanoma cells to shut down cell cycle machinery and adopt a stressed, hypertrophic phenotype. These findings redefine the kinetics of astrocyte-microglia/macrophage/cancer cell crosstalk in brain metastasis, uncovering the disruption of this interplay as a promising therapeutic approach for dismantling immune evasion in brain metastasis (17–19).

Selective TARA depletion impairs melanoma growth by 34-fold in late stages, reducing 3-fold that of early stages. Several preclinical models have previously shown a partial dependency for astrocytes for intracranial disease progression (11,15), although usually revealing more discrete phenotypes probably due to incomplete niche ablation or compensatory stromal mechanisms (20). We uncovered that TARA not merely represents a physical support or structural scaffolding through cell-cell interactions, such as Lgals1-Itgb1, Vim-Cd44, but establish an active, specialized metabolic niche for the growing tumor.

Spatial transcriptomics revealed that peritumoral niche not only provides physical support for cancer cell invasion, but primes energetic and biosynthetic demands of cancer cells under extreme microenvironmental stress. While some clues regarding metabolic hijacking have been found in glioblastoma matrisomes (21,22), a precise spatial description was missing regarding peritumoral TARA-cancer cell crosstalk that leads to tumor invasion and growth (23). Losing this metabolic niche may account for reduced protumoral cues in the TME (20), directly impairing cell cycle progression (24). Furthermore, this collapse rewires not only the immediate peritumoral space but the broader brain response towards cancer cells, resembling an intrinsic antitumor immune response (25).

The selectivity and efficiency of our target manipulation was confirmed by the post-depletion presence of star-like, resting state astrocytes close to the constrained foci, as close as 100 µm from the tumor border. This restricted effect supports the pathological role of invading cancer cells in corrupting and activating homeostatic astrocytes in close, direct contact with the invasive front (11,26). The lack of supporting peritumoral niche allowed the subsequent massive infiltration of Iba-1+ cells into the tumor mass. This is highly consistent with literature describing the physical isolation of brain tumors by reactive astrocytic scar (3,27). TARA secrete dense extracellular matrix components and soluble mediators that trap immune cells in the peritumoral scar, preventing them from contacting the tumor core (28). Thus, our results show that TARA depletion unleashes myeloid cells to infiltrate the invading foci and perform a robust antitumor inflammatory response, which is associated with local inflammation and interferon secretion, as reported conceptually by several studies (29,30). Moreover, proinflammatory microglia actively contribute to metastatic impairment through improved immune response (31). Indeed, upon spatial transcriptomics analysis, TARA-depleted tumors exhibited 100% positive spots for activated microglial signature compared to a mere 35% in control tumors. This striking result goes in line with previous reports indicating that infiltrating microglia and macrophages can act as required, highly potent antitumor cell subsets in brain cancer if properly activated (32). However, to our knowledge, no prior study has successfully linked the physical dismantling of the peritumoral astrocytic niche with an improved, unhindered innate antitumor response and subsequent enforcement of cancer cell tumorostatic phenotype.

Importantly, melanoma dependency on TARA is not restricted to late tumor growth but dictates early tumor development as well. Elimination of TARA from day 4 to 7 revealed a 3-fold reduced tumor burden, accompanied by iba-1^+^ cell influx. Intriguingly, immunohistochemical analysis showed that melanoma cells did not undergo programmed cell death (remaining Caspase-3 negative), but instead exhibiting a striking 3-fold increase in cellular and nuclear size. This morphological transformation in highly consistent with an Interferon and TNF-α mediated cell cycle arrest operating upon downstream p53 and p21 activation, congruent with well-stablished principles of cytokine-induced senescence (33–35). Consistently, this inflammation and interferon-mediated p53 and p21-driven cell senescence was confirmed in late stages, with dismantled cell cycle machinery coupled with the upregulation of strong cell-cycle brakes and the profound downregulation of E2F targets, G2M checkpoint regulators, Myc signalling and histone transcription, a transcriptomic signature described in interferon-driven senescent states across peripheral tumors (36,37). Moreover, tumorostatic arrest was actively driven by the antitumor Iba-1^+^ cells found infiltrating the tumor from the periphery up to the core, with antigen presenting potential and TNF-α pathway enrichment, via NF-κB. This interferon rich hostile TME may force surviving melanoma cells to switch on secondary adaptations to protect themselves from immune-mediated clearance (38,39). Intriguingly, our cell-cell communication analysis revealed that the Mif-CD74 and Apoe-Scarb1, along with Vim-CD44 and Lgals1-Itgb1 axes mediated myeloid cell-cancer cell interactions specifically within the TARA-depleted niches. Mif-CD74 interplay is well described a sophisticated tumor counter-mechanism used to swift microglia/macrophage phenotype from antitumor, pro-inflammatory towards protumoral, tolerogenic one (40). Concurrently, the upregulation of the Apoe-Scarb1 axis may represent a protecting adaptation. Apoe may act as a metabolic support for cancer cells to cope with deprived and hostile TME and a resisting mechanism for ferroptosis mediated oxidative stress(41,42). This critical finding may represent tumor celĺs last resort to build an immunological shield and impede complete elimination by immune cells (43).

Taken together, our results reveal astrocyte-dependent melanoma growth in the brain, sustained through aggressive TME remodeling, metabolic provisioning and immune shielding. Targeting TARA rewires brain metastatic microenvironment leading to pro-inflammatory, antitumor microglial and macrophage infiltration. This spatial and metabolic niche disruption forces cancer cells to adopt stressed, non-proliferative survival state, reprogramming their phenotype from proliferative to tumorostatic, resistant cells. Thus, combined therapeutic targeting of TARA alongside the inhibition of the compensatory Cd74-Mif and Apoe-Scarb1 interactions may impair tumor growth in the brain through separate, highly synergistic pathways, opening new therapeutic opportunities to enhance the efficacy of immune checkpoint inhibitors and targeted therapies in brain metastasis.

## Materials and methods

### Animals

Eight to ten week-old female GFAP-Tk mice (B6.Cg-Tg(Gfap-Tk)7.1Mvs/J, Jackson Laboratory, Bar Harbor, ME) were used for *in vivo* experiments in accordance with Institutional Animal Care and Use Committee Protocols. Transgenic mice (TG) express the herpes-simplex virus Thymidine kinase gene (HSV-Tk) driven by the mouse glial fibrillary acidic protein (GFAP) promoter. Wild type littermates were used as controls (WT).

### Cancer cell line

B16-F10 melanoma murine cell line (ATCC, LGC Standards SLU, USA) was used in this study. B16-F10 cells were maintained in DMEM supplemented with 10% fetal bovine serum (FBS), penicillin (100 U/ml), streptomycin (100 µg/ml) and amphotericin B (0.25 µg/ml) (Thermo Fisher Scientific, MA, United States). Cells were discarded after twelve passages and substituted by new batches.

### Melanoma brain tumor development

Murine B16-F10 melanoma cells were diluted in sterile PBS and 2 µL of the cell solution containing 2×10^4^ cells/animal was intracranially injected into the striatum by stereotactic surgery (coordinates from bregma: +1 mm anteroposterior, +1.5 mm mediolateral, and -2.5 mm dorsoventral) under deep isoflurane anesthesia. Injection was performed with a beveled 33-gauge Hamilton syringe coupled to a microinjection pump (World Precision Instruments) at a rate of 0.4 ul/min. After injection, the needle was left in place for 10 min to avoid cell leakage. Animals received intraperitoneal buprenorphine (0.1 mg/kg) during and after surgery to minimize pain and were kept under observation for 72 hs. Littermates were divided in 2 groups (n=3) based on their genotype and GCV treatment schedule (Supplementary Figure 1A). Animals were sacrificed after 7 days (for early tumor development) and 12 days of tumor growth. 12 days *In vivo* assay was carried out three times (n = 6/group total). Early tumor growth (7 days) was carried out once (n=3/group).

### *In vivo* selective killing of proliferating TARAs

Mice received i.p. injections of GCV (100 µg/g/day) (200 µl) or saline from day 4 to day 6 both included (early tumor development) and from day 4 to 11 for late tumor development. This treatment only targets GFAP-expressing proliferating cells, selectively killing only tumor-reactive proliferating astrocytes during brain metastatic colonization.

### Histochemical analysis

For tumor area quantification, 10 µm thick paraffin embedded brain sections were stained using H-E. 2 different brain sections were analyzed separated by 100 µm between each other. Inmunohistochemical analysis was carried out in paraffin-embedded brain tissue arrays incubated with sodium citrate antigen retrieval pH= 6 for 20 min at 95°C. Then, peroxidase activity and unspecific unions were blocked through 3% hydrogen peroxide (PanReac AppliChem) in PBS for 30min at room temperature, followed by 1h with 0.4% Triton X-100 and 10% FBS in PBS. The samples were incubated with anti-GFAP (1:200, MA5-12023.) from Invitrogen, anti-Iba-1 (1:200, Ref. ab86607) from Abcam, anti-Ki67 (1:25, Ref. 550274) from BD PharMingen, and anti-Caspase-3 (1:200, Ref. M0851) from Dako overnight at 4°C. Then, the samples were incubated for 1h with specific biotinylated secondary antibodies (Thermo Fisher Scientific): goat anti-rabbit (1:1,500, Ref.31823), rabbit anti-mouse (1:2,000, Ref. 31813) and goat antirat (1:500, Ref. 31830). Protein expression was detected incubating the samples with with streptavidin-horseradish peroxidase (HRP) conjugated (1:500, Ref. 434323) for 30 minutes and visualized using diaminobenzidine (DAB) substrate (Ref. 34002), both from Thermo Fisher Scientific. Finally, samples were counterstained with hematoxylin, mounted and analyzed through FIJI Image J.

### Cancer cell nuclei measurement

For nuclei size calculation, images were analyzed using FIJI Image J software. Individual cells were measured in each tumor and the average nuclei size was calculated. At least 80 nuclei were analyzed for each tumor in three different samples for early tumor growth and four different samples (n=4/group) for late tumor growth.

### Intratumoral cell infiltration measurement

The infiltration plots for Iba-1 and Gfap positive cells were carried out through ImageJ software. In detail, immunohistochemistry images for both markers of control and TARA-depleted tumors were opened and color channels were separated. Next, the expression of Iba-1 and Gfap were highlighted while reducing the noise of unspecific interference, to match the original expression pattern (black for Iba-1/Gfap expression and white for the background). Finally, the image color pattern was inverted, and the plots were generated.

### Visium data

Four coronal brain sections from four animals were captured on two Visium slides (VLP156_A1: control animals C1 and C2; VLP156_D1: TARA-Depleted animals 1 and 2). C1, C2 and TARA-Depleted 2 contain tumor; TARA-Depleted1 contains no tumor and was used only as a negative control in the interaction analysis. Raw spot-by-gene UMI counts from Space Ranger were used throughout. Spots were clustered in Seurat (SCTransform), and the clusters were grouped into neuron, striatum, melanoma, oligodendrocyte and transitional superclusters. These clusters were later refined with cell-type estimation to define compartments.

### Single-cell reference

A single-cell reference was assembled from public datasets. Astrocytes were obtained from GSE226207 (cortical stab-wound injury) and GSE148611 (Aldh1l1-eGFP astrocytes following saline or LPS treatment). Reactive and proliferating states were identified using gene programs derived from within-dataset pseudobulk differential expression, with sample as the replicate. Cells with programmed scores above the 99th percentile of control cells within each dataset were assigned to the corresponding state. The proliferating program included Mki67, Top2a, Cdk1 and Ccnb1 and was retained only in GSE226207 (532 cells). Microglial populations were obtained from three wild-type samples of GSE285670 (CD45⁺CX3CR1⁺ cells from Nras melanoma brain metastases), comprising inflammatory (1,068 cells), DAM-like (1,446) and proliferating microglia (1,233). RelA-knockout samples were excluded. Additional normal-brain populations were obtained from the Allen whole-mouse-brain atlas (10x v3), capped at 3,000 cells per population, and 1,341 melanoma cells were obtained from GSE156444.

### Cell-type estimation

Cell-type abundances were estimated using cell2location v0.1.5 with raw counts for 13,682 shared genes. Reference signatures were estimated by negative-binomial regression with sample as batch (250 epochs). Spatial mapping used section as batch, five expected cells per spot and detection alpha of 20 (10,000 full-batch epochs, 1,000 posterior samples, seed 0). Cell-type proportions were calculated by dividing each label’s posterior mean abundance by the total abundance per spot. To obtain estimated cell numbers, these proportions were multiplied by nuclei counts derived from H&E images using StarDist (2D_versatile_he model) through Spotiphy v0.3.0, with nuclei assigned to spots by their centers. As a sensitivity analysis, spatial mapping was repeated using the mean nuclei count per section as the expected number of cells per spot (C1: 11.17; C2: 8.09; TARA-depleted1: 4.96; TARA-depleted2: 7.53).

### Tissue compartments and glial abundance

Tumor boundaries were determined from cell2location-estimated melanoma proportions using an iterative, section-specific half-maximum threshold. Starting from 0.15, thresholds were updated as background + 0.5 × (plateau − background), where background was the median melanoma proportion in spots at least five neighbor steps from the tumor and plateau was the 75th percentile within the tumor. Retained tumor spots were required to have at least one tumor neighbor. Boundaries stabilized after 3–4 iterations, yielding thresholds of 0.250, 0.236 and 0.137 and tumor regions containing 813, 496 and 45 spots in C1, C2 and TARA-depleted2, respectively.

Spots within two neighbor steps outside the tumor were classified as striatal grey matter, cortical grey matter or white matter according to their Pearson correlation with same-section supercluster expression centroids. Centroids were constructed using the top 50 Wilcoxon marker genes per supercluster, excluding genes with higher mean expression in the tumor. Border analyses focused on the striatal peritumor region (70, 55 and 64 spots in C1, C2 and TARA-depleted2, respectively), as TARA-depleted lacked a cortical tumor border.

### Spatial ligand–receptor analysis

Ligand–receptor (LR) activity was inferred using stLearn v1.4.1 with library-size-normalized, log-transformed expression and literature-supported mouse LR pairs from connectomeDB2020. LR hotspots were identified by comparing expression across each spot and its immediate neighbors with 500 expression-matched background gene pairs (Benjamini–Hochberg-adjusted P ≤ 0.05).Spatial interactions were assessed at the tumor border for proliferating astrocytes and within the tumor and striatal peritumor for inflammatory microglia. An adjacent glia–melanoma spot pair was considered LR-active when at least one spot was an LR hotspot and both ligand and receptor expression were detected across the pair. Glial presence required at least one estimated cell, whereas melanoma presence required a deconvolved proportion greater than 0.15. Astrocyte LR pairs were ranked by the smaller border interaction count across the two controls, and microglial pairs by their interaction counts within the TARA-depleted tumor.

### Validation in available human dataset

Further validation was performed in a bigger patient cohort using publicly available data from R2: Genomics Analysis and Visualization Platform (http://r2.amc.nl http://r2platform.com), (Tumor Skin Cutaneous Melanoma - TCGA - 375 – rsem - tcgars: Source: TCGA ID: SKCM Date: 2014-01-21), The correlation between the expression of CD74, BST2, CD68 and CD86 and overall survival was analyzed in melanoma patients (data show median value for all the genes). For Cdkn1a correlation, R2 Genomics Analyst was used. In detail, negatively correlated genes in “biological process” were obtained and KEGG pathway in Enrichr model analysis was carried out. 3909 combinations meet the inclusion criteria, (R cutoff of 0 neg as R correlation p-value<=0.05, Multiple testing correction applied: False Discovery Rate). Results were limited to 1500 rows. Results show p value ranked top 10 pathways negatively correlated with Cdkn1a (p21).

### Statistics analysis

Data are expressed as mean ± SD. Statistical analyses of the in vivo data were performed using GraphPad Prism software. First, normality tests were performed to assess whether the data followed a normal distribution. Based on the resulting p value, Welch’s t-test was used to compare the control and TARA-treated brains. Statistical significance was set at p ≤ 0.05 for all comparisons.

## Supplementary Figures

**Supplementary Figure 1:**
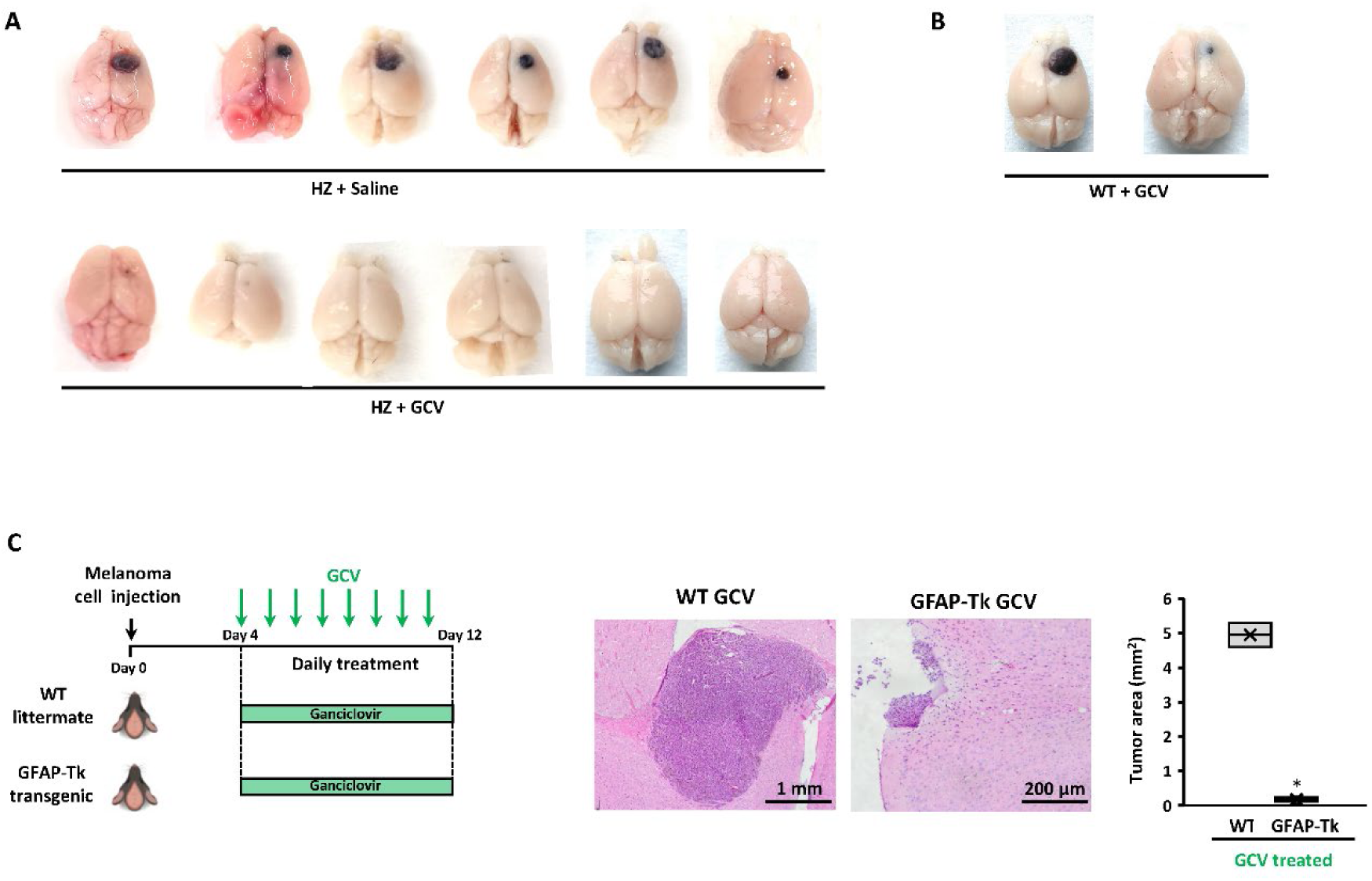
In vivo results from different assays. A) Macroscopic images from the brains of both experimental groups from all the in vivo assays carried out. B) Brains from WT mice treated with GCV for 12 days. C) Experimental design for the possible GCV-mediated antitumor effect in WT mice for control (n=2), describing methodology, treatment scheme and in the right panel the in vivo results. Differences between WT and TARA-depleted group were considered statistically significant for *p< 0,05.

**Supplementary Figure 2:**
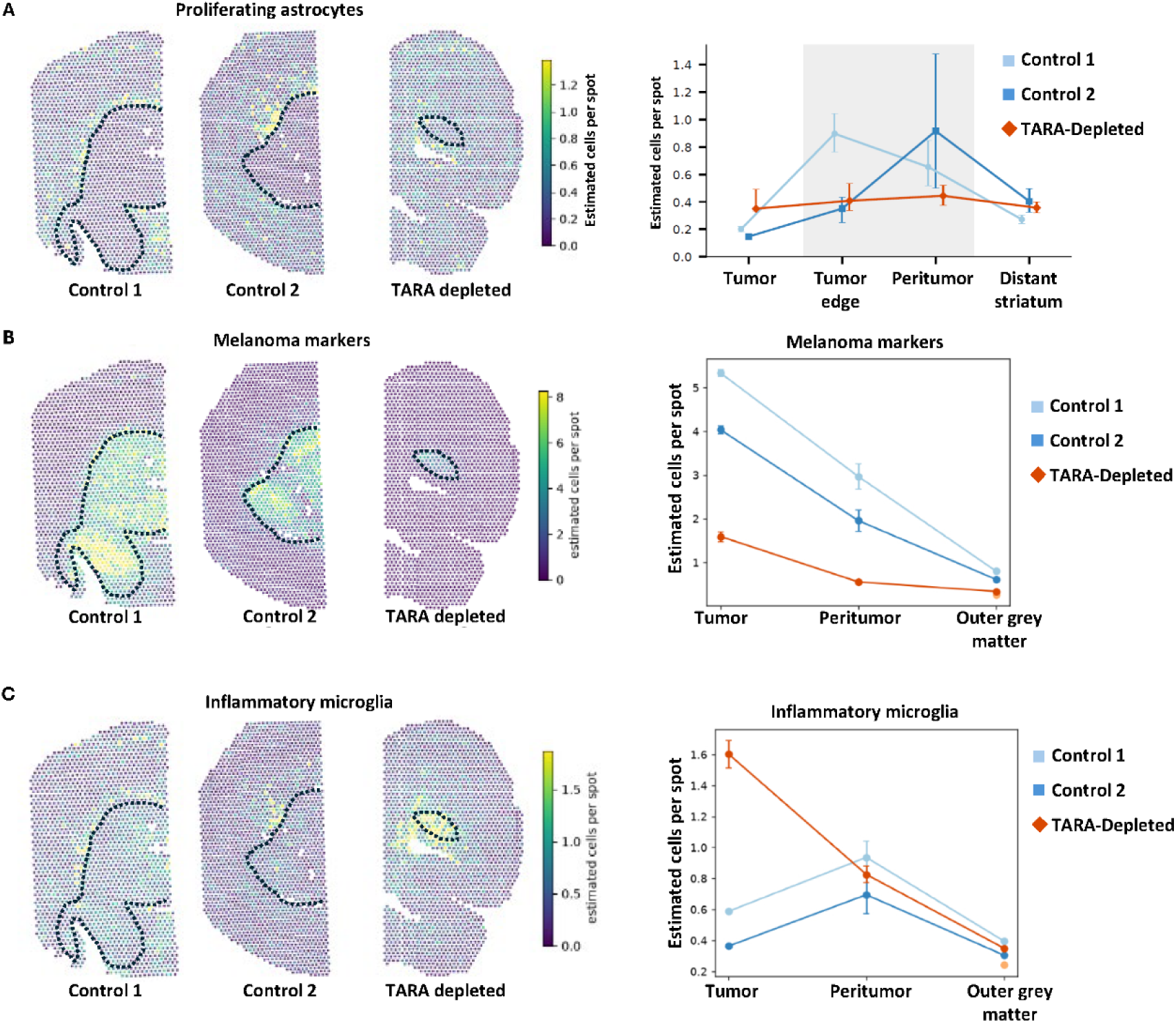
Spatial distribution of estimated cell number for melanoma and inflammatory microglia. A) Estimated melanoma cells per spot in the brain tumors, peritumoral niche and outer grey matter from Control and TARA-depleted mice brain. B) Estimated inflammatory microglia cells per spot in tumor, peritumoral niche and outer grey matter in tumors from Control and TARA-depleted mice.

**Supplementary Figure 3.**
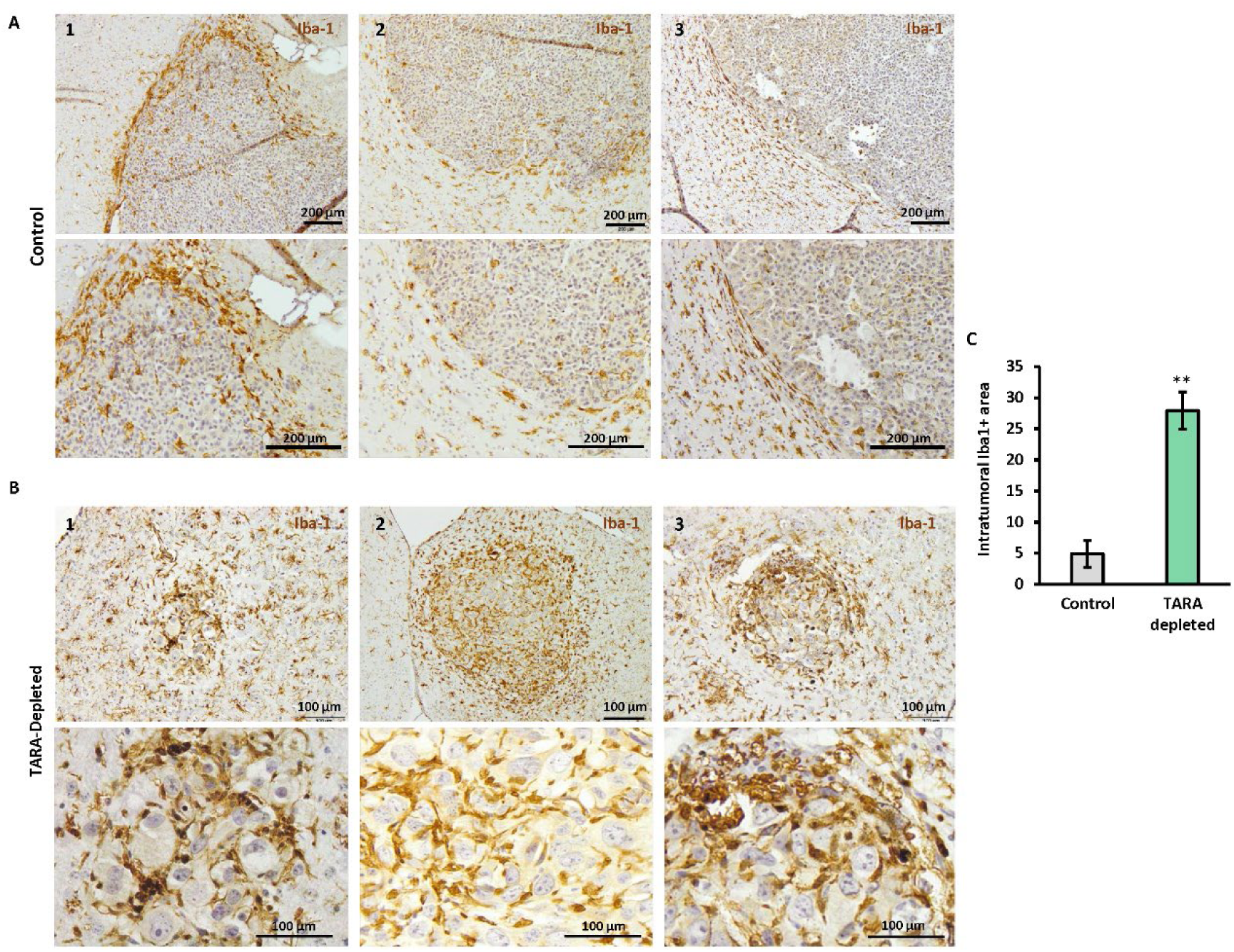
Representative images of Iba-1+ cell distribution in late tumor growth melanoma samples (12 days). A) Iba-1+ immunohistochemistry in control melanoma tumors showing peritumoral Iba-1^+^ cell distribution with mainly peripheral scarce infiltration. B) Iba-1 immunohistochemistry revealing Iba-1^+^ cell influx towards melanoma tumors, invading almost to the tumor core and surrounding hypertrophic cancer cells. Data were considered statistically significant for ** p< 0.01.

**Supplementary Figure 4:**
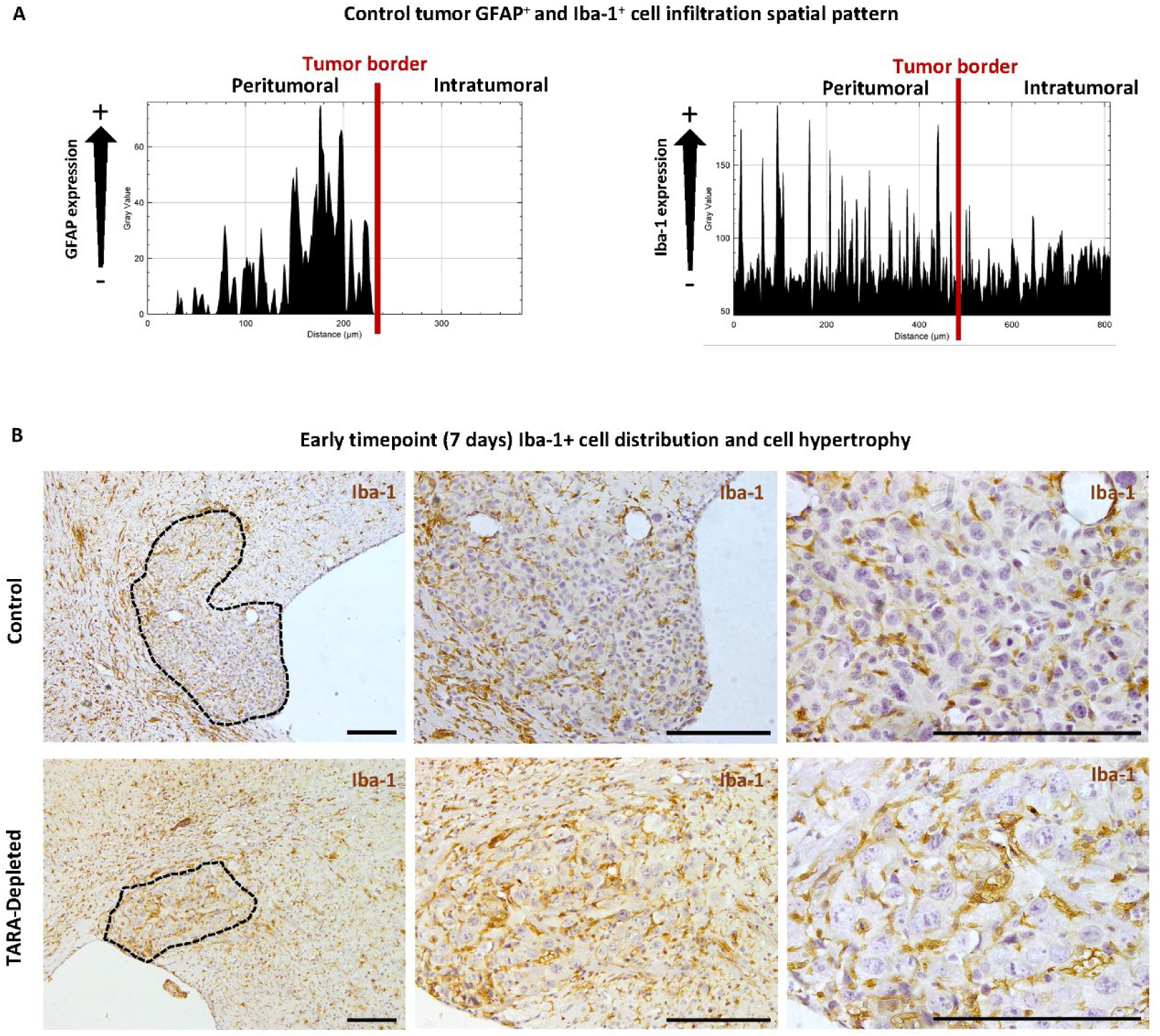
Spatial distribution of GFAP^+^ and IBA-1^+^ cells in the tumor microenvironment. A) Spatial distribution of GFAP^+^ and IBA-1 cells from the peritumoral niche into the tumor mass in control mice melanoma tumors. B) Representative images of Iba-1^+^ cells in melanoma tumors of control and TARA-depleted mice after 7 days of tumor growth (3 days of GCV treatment), with hypertrophic cells with enlarged nuclei surrounded by Iba-1^+^ cells visible in TARA-depleted tumor. Scale bar: 200 µm.

**Supplementary Figure 5:**
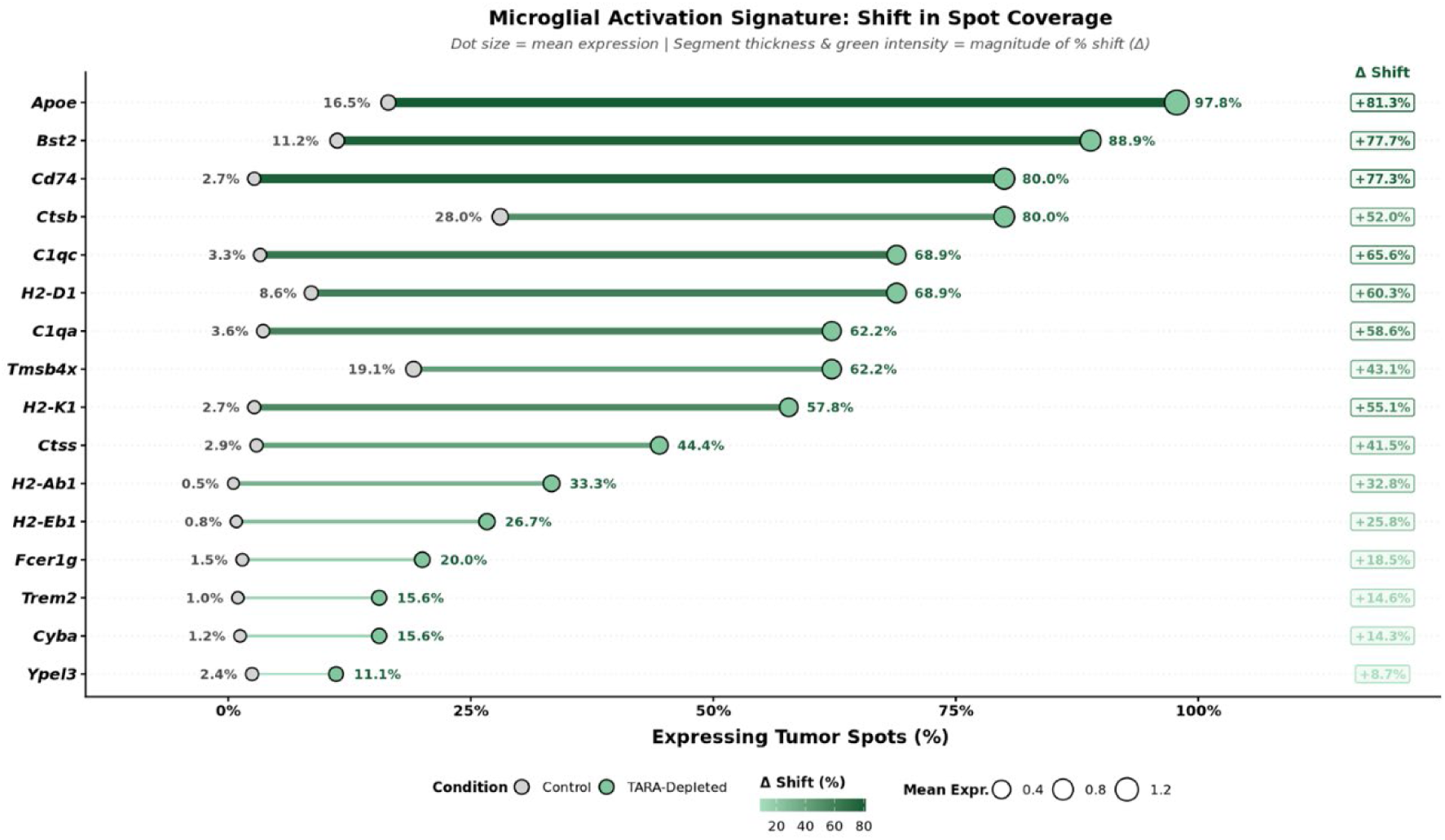
Microglial Activation Signature genes. Genes utilized to spatially map the activated microglia in melanoma tumors. The percentage of tumor niche spots positive for this transcript signature in represented using gray dot for control spots and green dot for TARA-depleted spots. The increase in abundance is represented in the squares in the right margin.

**Supplementary Figure 6:**
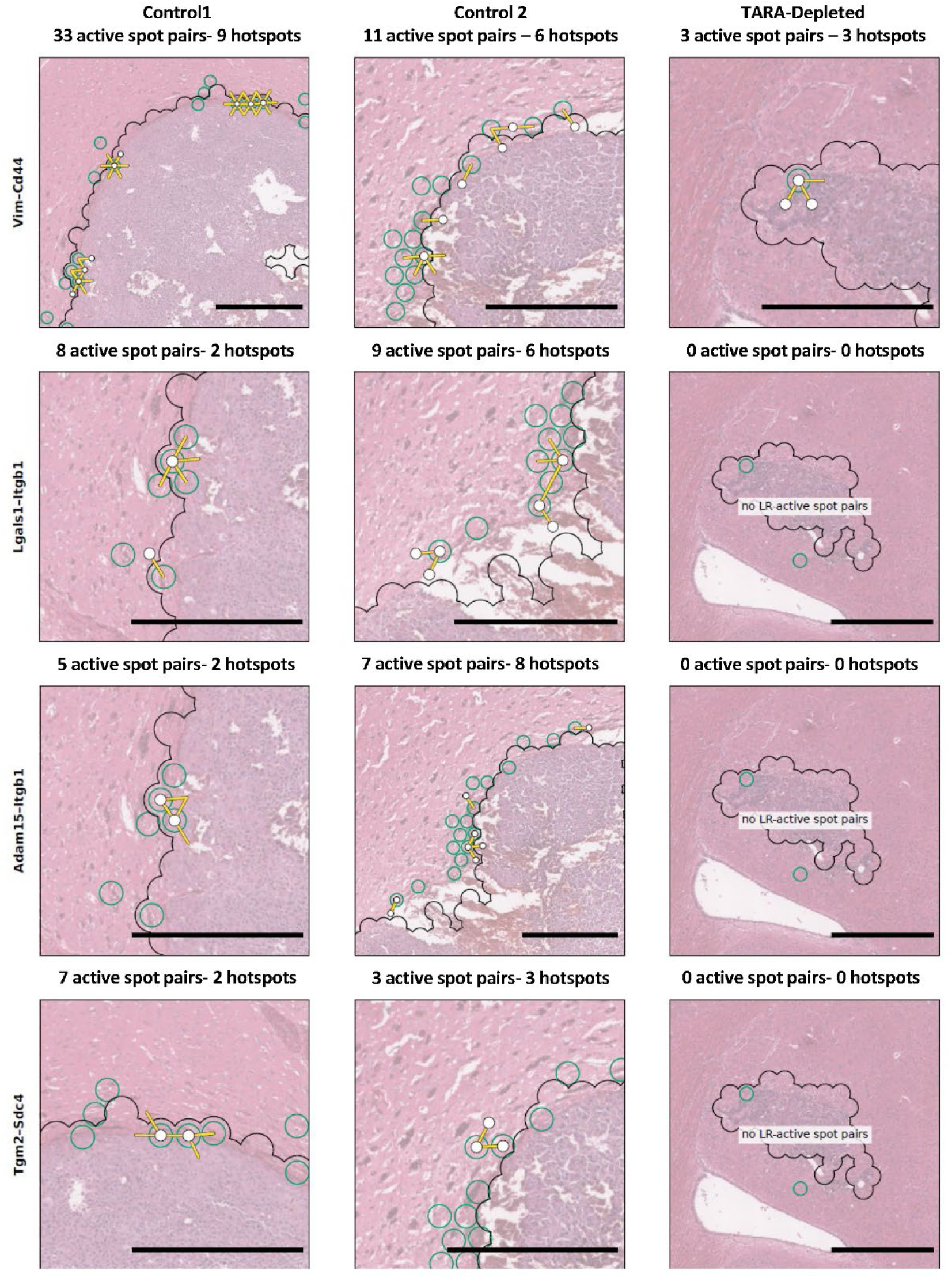
Cell-Cell interaction in the tumor border between proliferating astrocytes and melanoma cells. Images showing L-R pairs occurring during the crosstalk between proliferating astrocytes (green circles) and melanoma cells (area inside the black line) regarding Vim-Cd44 (first row), Lgals1-Itgb1 (second row), Adam15-Itgb1 (third row) and Tgm2-Sdc4 (last row). Scale bar 500 µm.

**Supplementary Figure 7:**
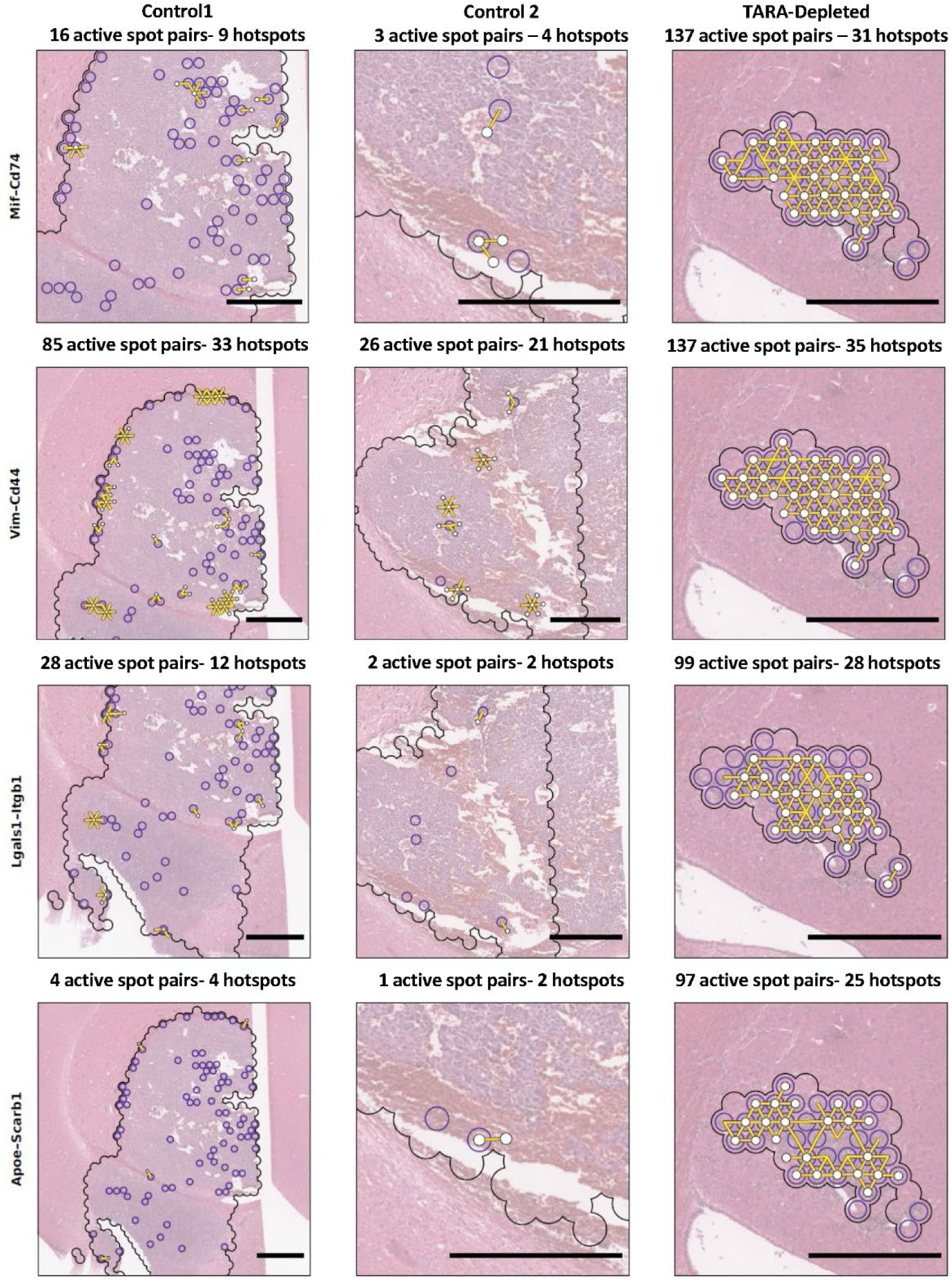
Cell-Cell interaction in the tumor core between inflammatory microglia/macrophages and melanoma cells. Images showing L-R pairs occurring during the crosstalk between inflammatory microglia/macrophages (purple circles) and melanoma cells (area inside the black line) regarding Mif-Cd74 (first row), Vim-Cd44 (second row), Lgals1-Itgb1 (third row) and Apoe-Scarb1 (last row). Scale bar 500 µm.

## Author information

These authors contributed equally: Alba Herrero and Yibi Chen.

## Contributions

FN.S., B.A. and A.B. carried out transgenic mouse melanoma model. A.H. conducted immunohistochemistry and analyzed the data. Y.C. analyzed spatial transcriptomic data and generated new data. E.A. conducted immunohistochemistry assays. T.H. analyzed spatial transcriptomic data. A.X. performed Visium experiments. Q.N. provided expertise in spatial transcriptomics. A.B. wrote the manuscript. A.H., B.A., F.S. and Q.N. edited the manuscript. M.D.B, B.A. and A.B obtained funding. All authors read and approved the final version of the manuscript. A.B. conceived the study and supervised the work.

## Ethics declarations

All animal procedures were approved by The University of the Basque Country (reference 2021/53) and conducted in concordance with international guidelines.

## Data availability

The spatial transcriptomics data generated in this study will be deposited in the Gene Expression Omnibus (GEO) during peer review.

Publicly available single-cell RNA-sequencing datasets used for cell-type deconvolution were obtained from GEO under accession numbers GSE226207, GSE148611, GSE285670 and GSE156444, and from the Allen Institute whole-mouse-brain cell atlas (10x v3).

Additional data supporting the findings of this study are available from the corresponding author upon reasonable request.

## Acknowledgements

The authors would like to thank Maria Jesus Fernandez and Yolanda Martin for the technical support. The authors thank for technical and human support provided by SGIKER. The authors also acknowledge the University of Queensland Research Computing Centre for providing access to the Bunya high-performance computing infrastructure and computational support. A.B. would like to thank University of Queensland and QIMR Berghofer for hosting the research stay and proving access to its facilities.

