## Supplementary Figures for "Astrocyte-mediated immune shielding protects melanoma brain metastases from microglia/macrophage induced Interferon stress and metabolic collapse"

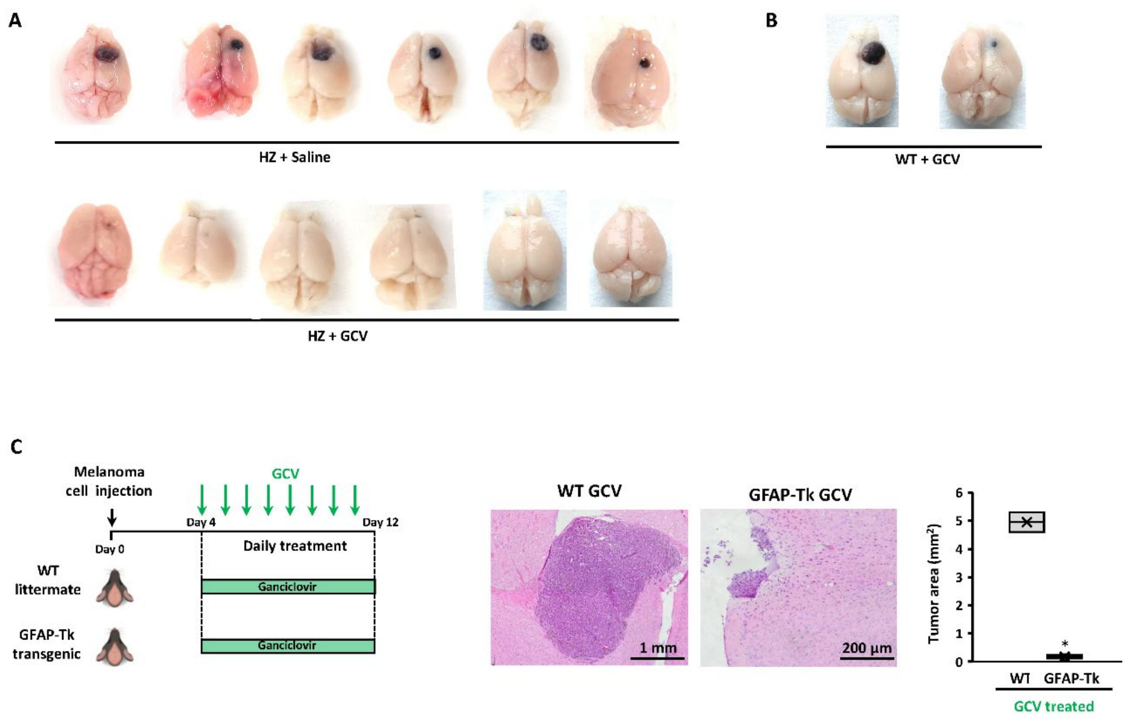


**Supplementary Figure 1**: **In vivo results from different assays.** A) Macroscopic images from the brains of both experimental groups from all the in vivo assays carried out. B) Brains from WT mice treated with GCV for 12 days. C) Experimental design for the possible GCV-mediated antitumor effect in WT mice for control (n=2), describing methodology, treatment scheme and in the right panel the in vivo results. Differences between WT and TARA-depleted group were considered statistically significant for *p< 0,05.

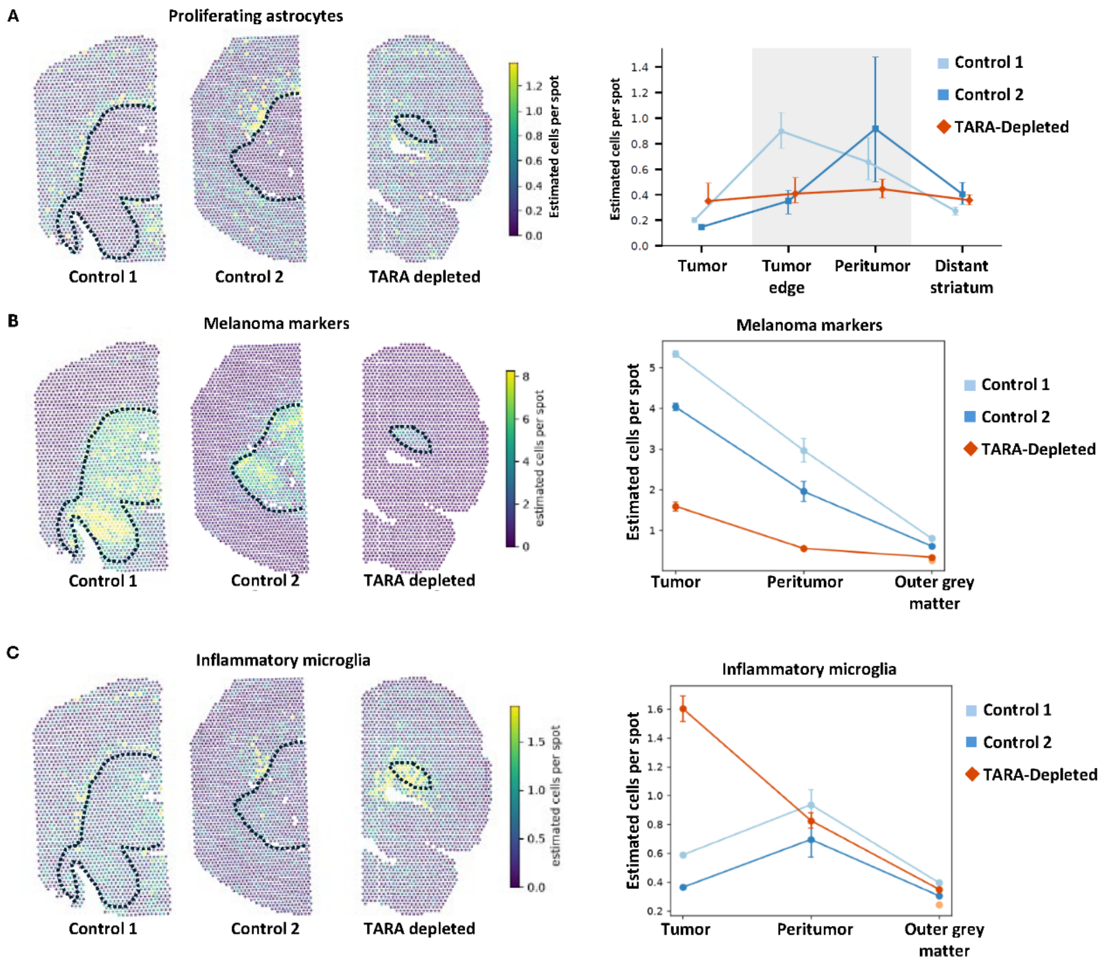


**Supplementary Figure 2: Spatial distribution of estimated cell number for melanoma and inflammatory microglia.** A) Estimated melanoma cells per spot in the brain tumors, peritumoral niche and outer grey matter from Control and TARA-depleted mice brain. B) Estimated inflammatory microglia cells per spot in tumor, peritumoral niche and outer grey matter in tumors from Control and TARA-depleted mice.

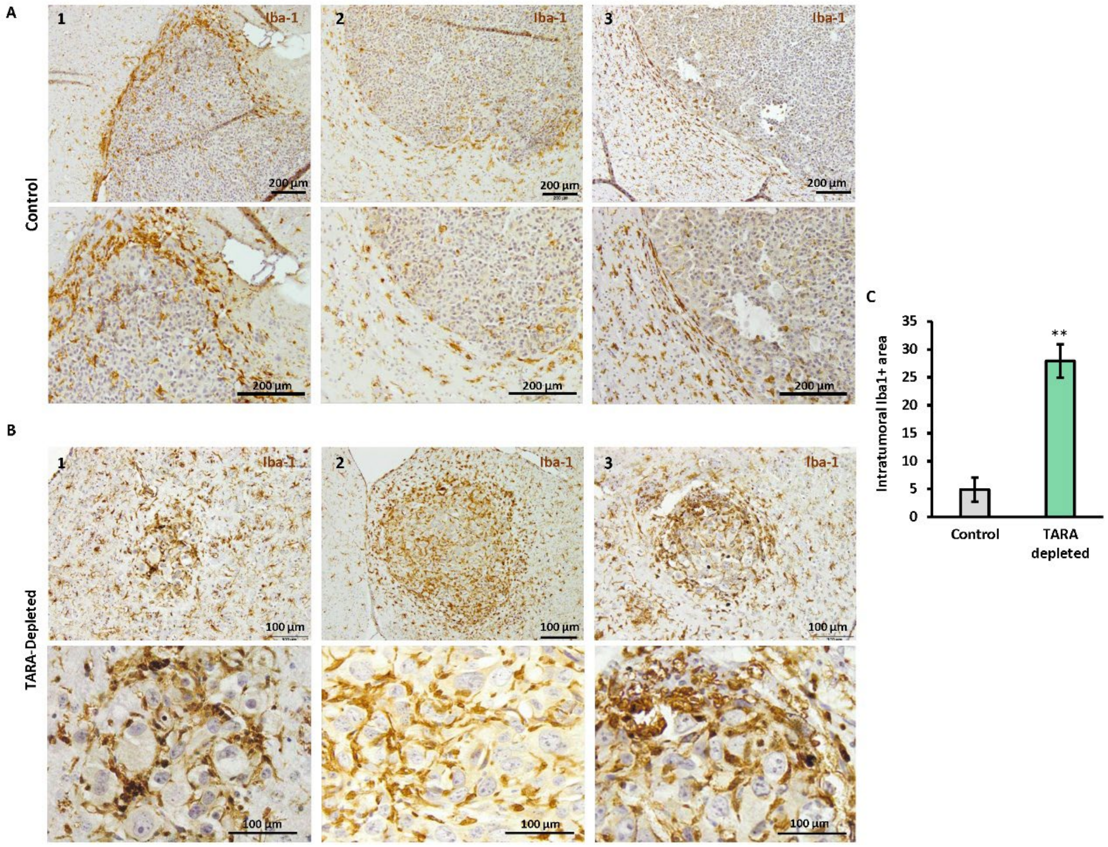


**Supplementary Figure 3. Representative images of Iba-1+ cell distribution in late tumor growth melanoma samples (12 days).** A) Iba-1+ immunohistochemistry in control melanoma tumors showing peritumoral Iba-1$^{+}$ cell distribution with mainly peripheral scarce infiltration. B) Iba-1 immunohistochemistry revealing Iba-1$^{+}$ cell influx towards melanoma tumors, invading almost to the tumor core and surrounding hypertrophic cancer cells. Data were considered statistically significant for ** $p< 0.01$.

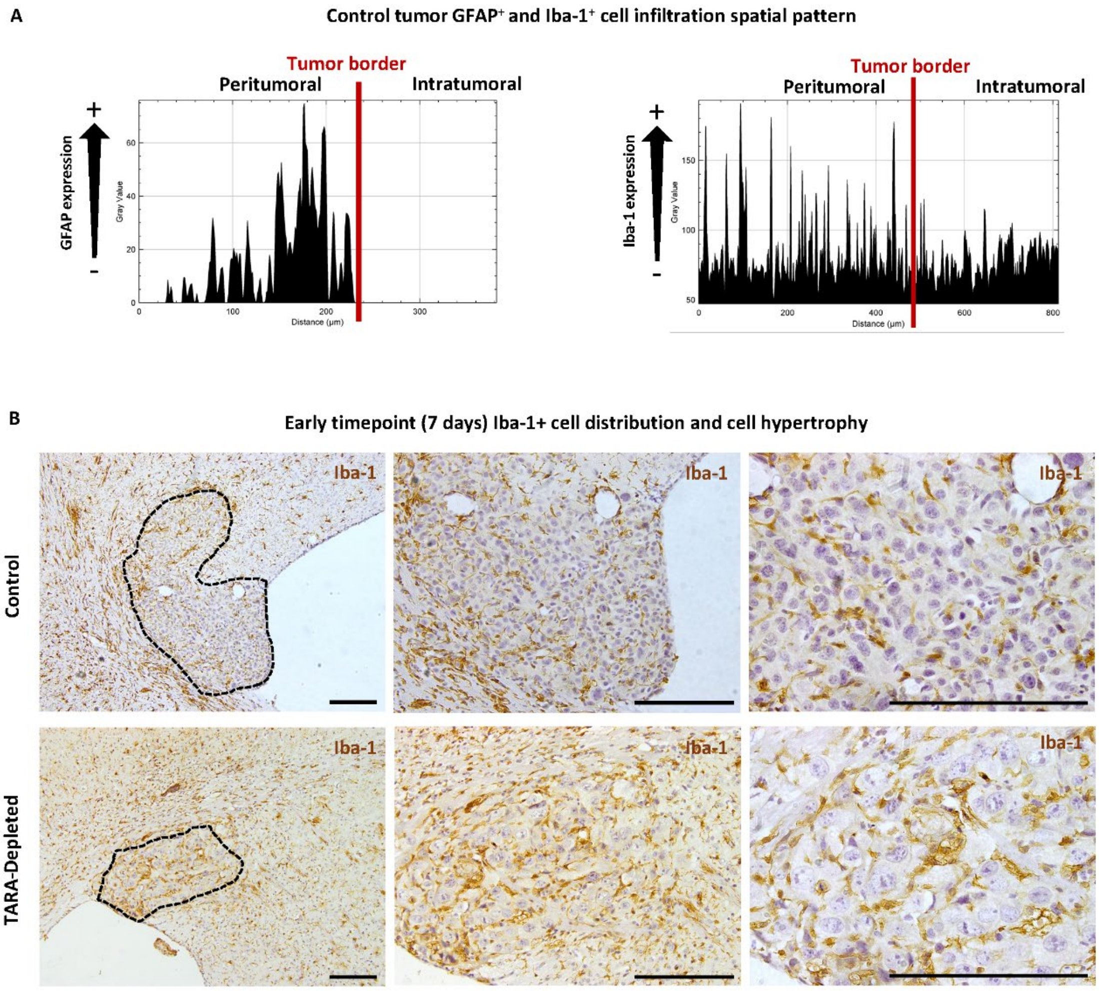


**Supplementary Figure 4: Spatial distribution of GFAP⁺ and IBA-1⁺ cells in the tumor microenvironment.** A) Spatial distribution of GFAP⁺ and IBA-1 cells from the peritumoral niche into the tumor mass in control mice melanoma tumors. B) Representative images of Iba-1⁺ cells in melanoma tumors of control and TARA-depleted mice after 7 days of tumor growth (3 days of GCV treatment), with hypertrophic cells with enlarged nuclei surrounded by Iba-1⁺ cells visible in TARA-depleted tumor. Scale bar: 200 µm.

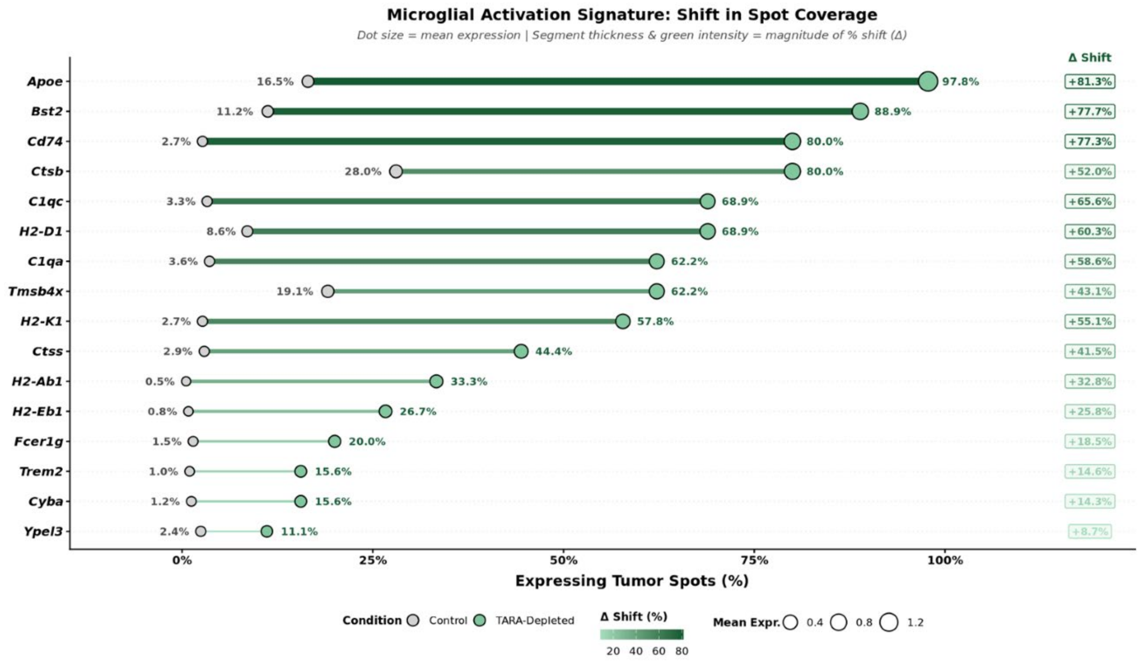


**Supplementary Figure 5: Microglial Activation Signature genes.** Genes utilized to spatially map the activated microglia in melanoma tumors. The percentage of tumor niche spots positive for this transcript signature in represented using gray dot for control spots and green dot for TARA-depleted spots. The increase in abundance is represented in the squares in the right margin.

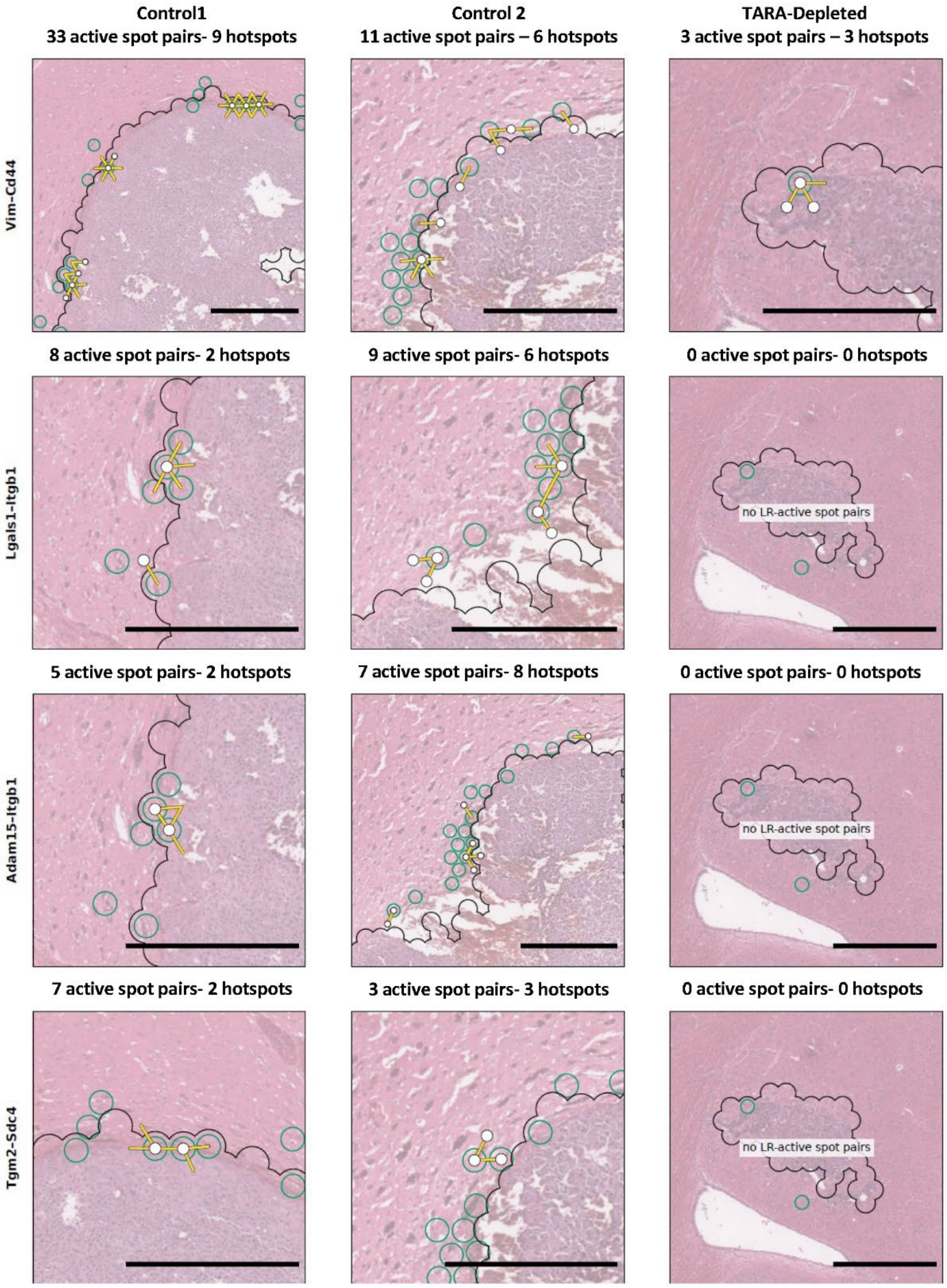


**Supplementary Figure 6: Cell-Cell interaction in the tumor border between proliferating astrocytes and melanoma cells.** Images showing L-R pairs occurring during the crosstalk between proliferating astrocytes (green circles) and melanoma cells (area inside the black line) regarding Vim-Cd44 (first row), Lgals1-Itgb1 (second row), Adam15-Itgb1 (third row) and Tgm2-Sdc4 (last row). Scale bar 500 µm.

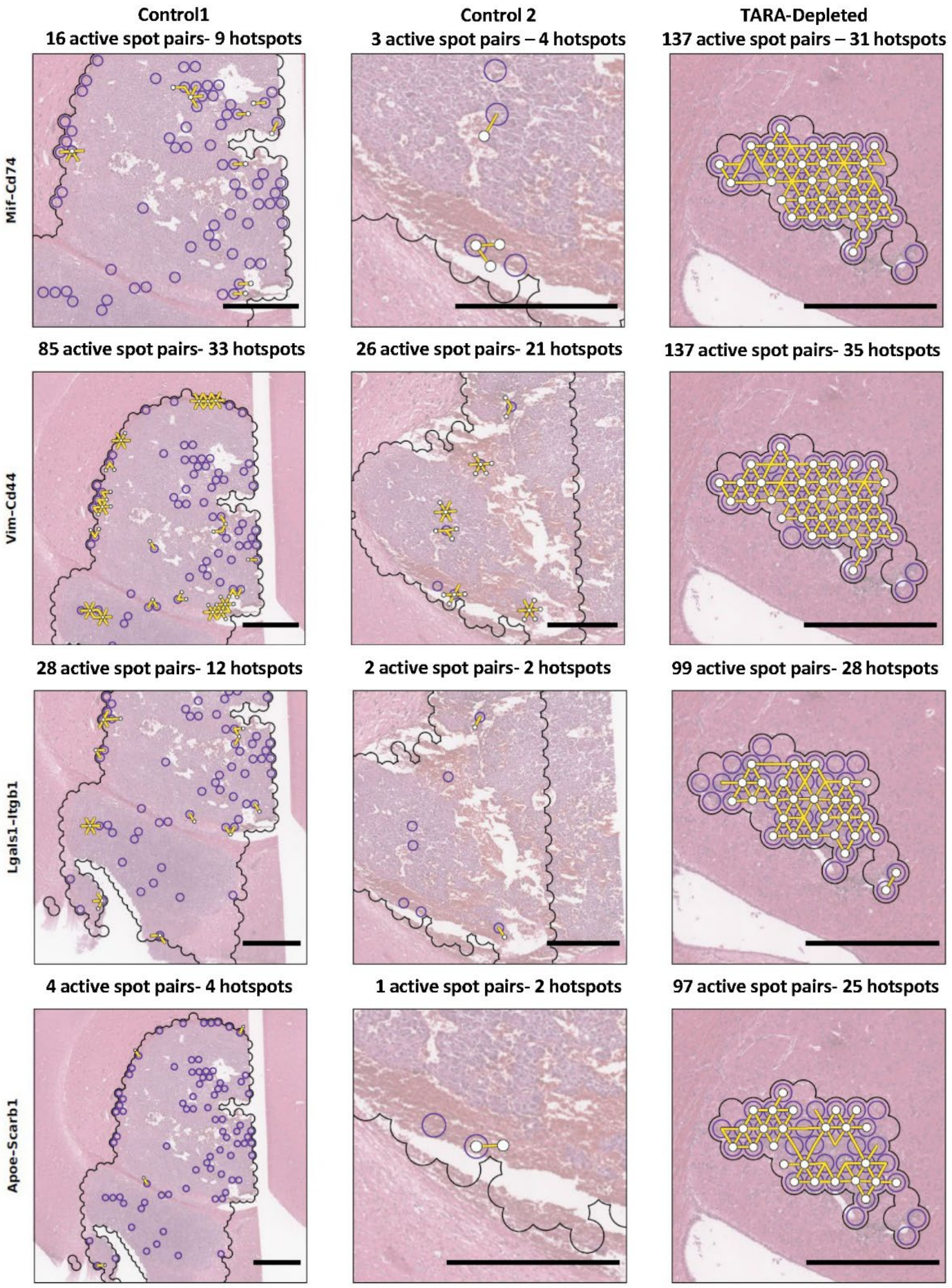


**Supplementary Figure 7: Cell-Cell interaction in the tumor core between inflammatory microglia/macrophages and melanoma cells.** Images showing L-R pairs occurring during the crosstalk between inflammatory microglia/macrophages (purple circles) and melanoma cells (area inside the black line) regarding Mif-Cd74 (first row), Vim-Cd44 (second row), Lgals1-Itgb1 (third row) and Apoe-Scarb1 (last row). Scale bar 500 µm.

**Funding:**

This work was supported by K2024/00003 and GV IT1524-22 from The Basque Government, GIU20/035 from The University of The Basque Country and research grant to A.B. from Fundación Jesus de Gangoiti Barrera. A.B. received a mobility grant (MV_2025_1_0029) for international research stay from the Basque Government.

**Author information:**

These authors contributed equally: Alba Herrero and Yibi Chen.

**Contributions:**

FN.S., B.A. and A.B. carried out transgenic mouse melanoma model. A.H. conducted immunohistochemistry and analyzed the data. Y.C. analyzed spatial transcriptomic data and generated new data. E.A. conducted immunohistochemistry assays. T.H. analyzed spatial transcriptomic data. A.X. performed Visium experiments. Q.N. provided expertise in spatial transcriptomics. A.B. wrote the manuscript. A.H., B.A., F.S. and Q.N. edited the manuscript. M.D.B, B.A. and A.B obtained funding. All authors read and approved the final version of the manuscript. A.B. conceived the study and supervised the work.

**Ethics declarations**

All animal procedures were approved by The University of the Basque Country (reference 2021/53) and conducted in concordance with international guidelines.

**Competing interest**

The authors declare no competing interest.

**Data availability**

The spatial transcriptomics data generated in this study will be deposited in the Gene Expression Omnibus (GEO) during peer review.

Publicly available single-cell RNA-sequencing datasets used for cell-type deconvolution were obtained from GEO under accession numbers GSE226207, GSE148611, GSE285670 and GSE156444, and from the Allen Institute whole-mouse-brain cell atlas (10x v3).

Additional data supporting the findings of this study are available from the corresponding author upon reasonable request.

**Acknowledgements**

The authors would like to thank Maria Jesus Fernandez and Yolanda Martin for the technical support. The authors thank for technical and human support provided by SGIKER. The authors also acknowledge the University of Queensland Research Computing Centre for providing access to the Bunya high-performance computing infrastructure and computational support. A.B. would like to thank University of Queensland and QIMR Berghofer for hosting the research stay and proving access to its facilities.
